# Efficient GNE myopathy disease modeling with mutation specific phenotypes in human pluripotent stem cells by base editors

**DOI:** 10.1101/2020.11.25.397711

**Authors:** Ju-Chan Park, Jumee Kim, Hyun-Ki Jang, Seung-Yeon Lee, Keun-Tae Kim, Seokwoo Park, Hyun Sik Lee, Hee-Jung Choi, Soon-Jung Park, Sung-Hwan Moon, Sangsu Bae, Hyuk-Jin Cha

## Abstract

Despite the great potential of disease modeling with the isogenic pairs of human pluripotent stem cells (hPSCs), the extremely low efficiency of precise gene editing in hPSCs remains a technical hurdle for this approach. Herein, we took advantage of currently available base editors (BEs) to epitomize the isogenic disease model from hPSCs. Using this method, we established 14 hPSCs that harbor point mutations on the *GNE* gene, including four different mutations found in GNE myopathy patients. Due to lesser activation of p53 by BEs than Cas9, a higher editing efficiency with BEs was achieved. Four different mutations in the epimerase or kinase domains of *GNE* revealed mutation-specific hyposialylation, which was closely correlated to pathological clinical phenotypes. These mutation-specific hyposialylation patterns were evident in GNE protein structure modeling. Furthermore, treatment with a drug candidate currently under clinical trials showed a mutation-specific drug response in GNE myopathy disease models. These data suggest that isogenic disease models from hPSCs using BEs could serve as a useful tool for mimicking the pathophysiology of GNE myopathy and for predicting drug responses.

## Introduction

Since the establishment of induced pluripotent stem cells (iPSCs), great advances in both autologous stem cell therapy and disease modeling from patient-specific iPSCs has been achieved (Rowe and Daley, 2019). The “disease-in-a-dish” model (Tiscornia et al., 2011) enables the identification of unknown disease mechanisms and drug screening once the disease phenotype indicator has been determined (Shi et al., 2017). However, the phenotypes arising from the disparate genetic backgrounds of the control and disease iPSCs are often more robust than those of the disease itself, which hinders translationally relevant comparison (Merkle and Eggan, 2013). The establishment of isogenic pairs from disease to control human pluripotent stem cells (hPSCs) or vice versa using precise gene editing technology was proposed as a way to minimize this bias (Merkle and Eggan, 2013; Musunuru, 2013).

Considering that the majority (58%) of genetic diseases result from point mutations (Rees and Liu, 2018), precise gene editing technologies such as base substitution based on homology-directed repair with Cas9 (hereafter referred to as HDR) (Izmiryan et al., 2018; Jackow et al., 2019) and base editing (BE) (Osborn et al., 2020) have been applied in human iPSCs to correct genetic mutations in patient iPSCs. However, it is generally accepted that the gene editing efficiency of Cas9 *via* HDR in hPSCs is extremely low compared to other cell lines (Mali et al., 2013), which remains a technical challenge for hPSC-based disease modeling. Thus, various approaches have been developed to improve editing efficiency (Kim et al., 2020; Maurissen and Woltjen, 2020). One recent study demonstrated that p53-dependent cell death, induced by a DNA double-strand break (DSB) that results from the endonuclease activity of Cas9, is responsible for its low gene editing efficiency in hPSCs (Ihry et al., 2018).

While HDR operates in tandem with DSB, base substitution by base editors (BEs) is achieved through deamination of the target base by deaminase conjugated with nickase Cas9 (nCas9), without inducing DSB (Gaudelli et al., 2017; Komor et al., 2016; Rees and Liu, 2018). Target sequence accessibility (hereafter referred to as “target-ability”) of BEs is determined by the protospacer adjacent motif (PAM) sequence that contains the canonical trinucleotide NGG (NGG-PAM). As the target-ability of BEs is restricted by the PAM sequence, BEs based on the NG-PAM sequence were recently developed to overcome this limitation (Huang et al., 2019; Nishimasu et al., 2018).

GNE myopathy (OMIM #605820), a rare autosomal recessive degenerative skeletal muscle disorder, is caused by bi-allelic loss of function (LOF) mutations of the *GNE* gene. The enzymatic activity of the GNE protein depends in turn upon N-acetylmannosamine (ManNAc) kinase activity and the epimerase activity of UDP-N-acetylglucosamine, which is the rate-limiting step in sialic acid biosynthesis. The reduced overall enzymatic activity of the GNE protein leads to hyposialylation, which is thought to cause the progressive muscle weakness characteristic of GNE myopathy(Noguchi et al., 2004). There are no FDA-approved therapies for GNE myopathy (Nishino et al., 2015). The only drug candidate in clinical trials is N-acetylmannosamine (ManNAc), the intermediate product in the synthesis of sialic acid (Xu et al., 2017). Due to the unavailability of human GNE disease muscle cells, immortalized cell lines (Pham et al., 2017) or mouse models (Chan et al., 2017) were alternatively used to study the mechanism. However, these models are insufficient to mimic pathophysiology in human muscle cells (Perlman, 2016). Thus, an appropriate “GNE myopathy in a dish” model is required to characterize human GNE pathophysiology.

To establish the isogenic disease model, we operationally defined the GNE disease model as attenuated sialic acid production in GNE mutants, to serve as a pathophenotype indicator. First, we demonstrated that BEs (Huang et al., 2019) were more efficient than HDR in hPSCs due to lesser induction of p53-dependent cell death. Using this method, we produced 14 hPSCs with point mutations on the *GNE* gene. Isogenic pairs of hESCs with four different mutations and myocytes derived from these mutant hESCs revealed mutation-specific hyposialylation. Additionally, the 3D structure modeling of each GNE mutant was consistent with altered GNE enzymatic activity, correlating with different degrees of hyposialylation. The differing responses of each mutant upon drug candidate treatment suggests that the mutation status of GNE patients is an important factor to consider in clinical trials of GNE myopathy drug candidates.

## Results

### Target-ability of point mutations causing GNE myopathy by base editors

BEs, such as the adenine base editor (ABE) and cytosine base editor (CBE), substitute the target base through the action of the deaminases TadA or APOBEC1, respectively, conjugated with nCas9, which induces single-strand breaks (SSB) (Fig. 1A). Both BEs and HDR are potential strategies for the production of isogenic pairs of disease models with hPSCs. Thus, prior to producing hPSCs for GNE myopathy modeling, we first investigated the “target-ability” of mutations causing GNE myopathy from the ClinVar database (https://www.ncbi.nlm.nih.gov/clinvar/) (Landrum et al., 2016). Among the total 63 mutations found in GNE patients that are categorized as “pathogenic” or “likely pathogenic,” 34 mutations (54%) occur by a transition point mutation (purine to purine or pyrimidine to pyrimidine) and are targetable with ABE or CBE (Fig. 1B and Table S1). Eight mutations (12.7%) are transversion mutations (purine to pyrimidine or vice versa), which are targetable with either the prime editor (PE) or HDR. Among transition point mutations, for establishing isogenic pairs for a GNE model (from mutant to wild-type or vice versa), 73.5% of wild-type (25 out of 34 mutations) (Fig. 1C) and 70.6% of mutants (24 out of 34 mutations) (Fig. 1D) are targetable by BEs. As mentioned above, 26.5% of wild-type (nine mutations) and 29.4% of mutants (10 mutations) cannot be edited due to the lack of a PAM sequence. The target-ability of BEs has greatly expanded due to advances in BE techniques using NG-PAM (Huang et al., 2019; Nishimasu et al., 2018) (e.g., NG-ABE or NG-CBE) (Figs. 1C and D). Interestingly, the relatively frequent spontaneous deamination of cytosine to uracil in nature may account for the differential utility of CBE (for C to T mutation) and ABE (for T to C mutation) for producing mutants and wild-type lines, respectively (Figs. 1C and D).

**Figure 1.**
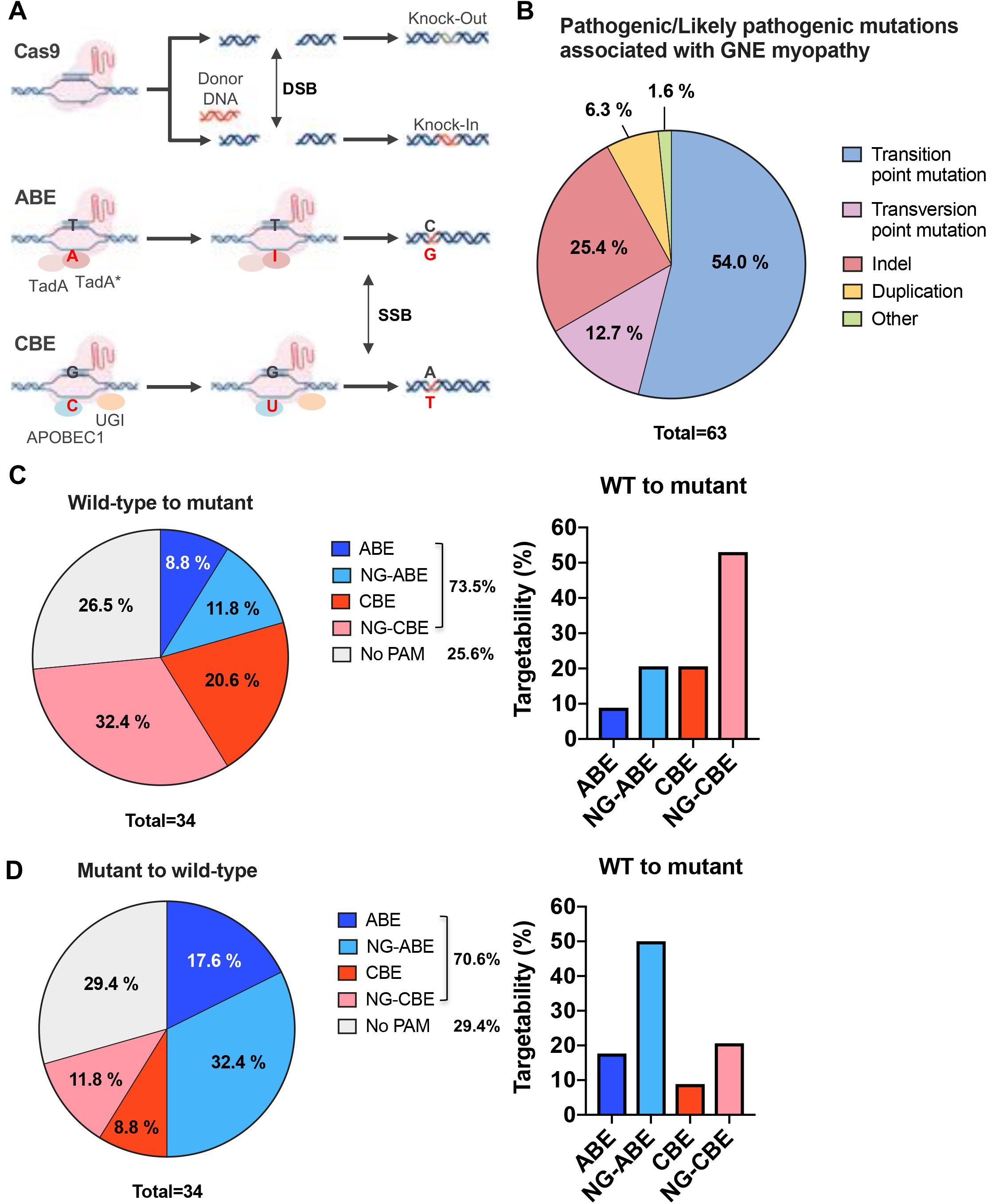
Target-ability of point mutations causing GNE myopathy by base editors. (A) Graphical description of three different gene editing technologies for generating point mutations (APOBEC1: cytidine deaminase, UGI: Uracil-DNA glycosylase inhibitor, TadA: adenine deaminase, TadA*: evolved TadA) (B) Pie Chart of distribution of GNE myopathy associated mutations descripted as pathogenic or likely pathogenic mutations of 63 patients in ClinVar database (https://www.ncbi.nlm.nih.gov/clinvar/) (C and D) Pie chart of targetability of mutations for ‘wild-type (WT) to mutant’ (C) or ‘mutant to wild-type (WT)’ (D) modeling by base editors (left), graphical presentation of target-ability of each base editor (right)

### Efficiency of base editing compared to HDR in hESCs

Considering that GNE myopathy occurs by loss of function (LOF) mutations of either the epimerase or kinase domains (Celeste et al., 2014), it is necessary to examine the functional effect of mutation in these domains of the GNE protein. For this, we chose four mutations of the epimerase (R160Q and I329T) and kinase (I588T and V272M) domains that would be readily achievable by ABE, CBE, and HDR (Fig. 2A). Compared to BEs, the HDR technique is more broadly targetable to all point mutations regardless of whether they are transversion or transition mutations (Rees and Liu, 2018). Thus, we first compared the editing efficiencies of all currently available techniques, HDR (Yang et al., 2013), ABE (ABEmax, (Koblan et al., 2018)), NG-ABE (NG-ABEmax, (Huang et al., 2019)), CBE (BE4max, (Koblan et al., 2018)) and NG-CBE (NG-BE4max, (Huang et al., 2019)), to generate point mutations of target sequences in hESCs. The efficiency of base substitution for R160Q by CBE was close to 37% (Fig. 2B), while those of NG-CBE and HDR were 4.1% and 0.93%, respectively (Figs. 2B and S1A). Similarly, 32.5% base substitution was achieved by ABE for I588T, while NG-ABE yielded 11.2% (Fig. S1B) and HDR yielded only 1% (Fig. 2C) substitution. For the V727M mutation, on-target base substitution efficiencies were 55.8% and 10.1% with CBE and NG-CBE (Fig. 2D and S1C), respectively, while HDR achieved 3.0% editing efficiency (Fig. 2D). Base substitution for I329T with HDR, ABE, and NG-ABE showed similar results, wherein the efficiency of HDR was lower (4.5% for I329T) than that of ABE (8.6%) and NG-ABE (6.3%) (Figs. 2E and S1D). However, the existence of bystander sequences in the target window reduced the observed percentage of base substitution without bystander mutation in V727M (Fig. S1E). Additionally, two mutants with bystander mutation (I329I and I329T) occurred in the I329T target due to codon degeneracy (Fig. S1F). The GNE mutant hESCs with bystander mutations are listed in Table S2.

**Figure 2.**
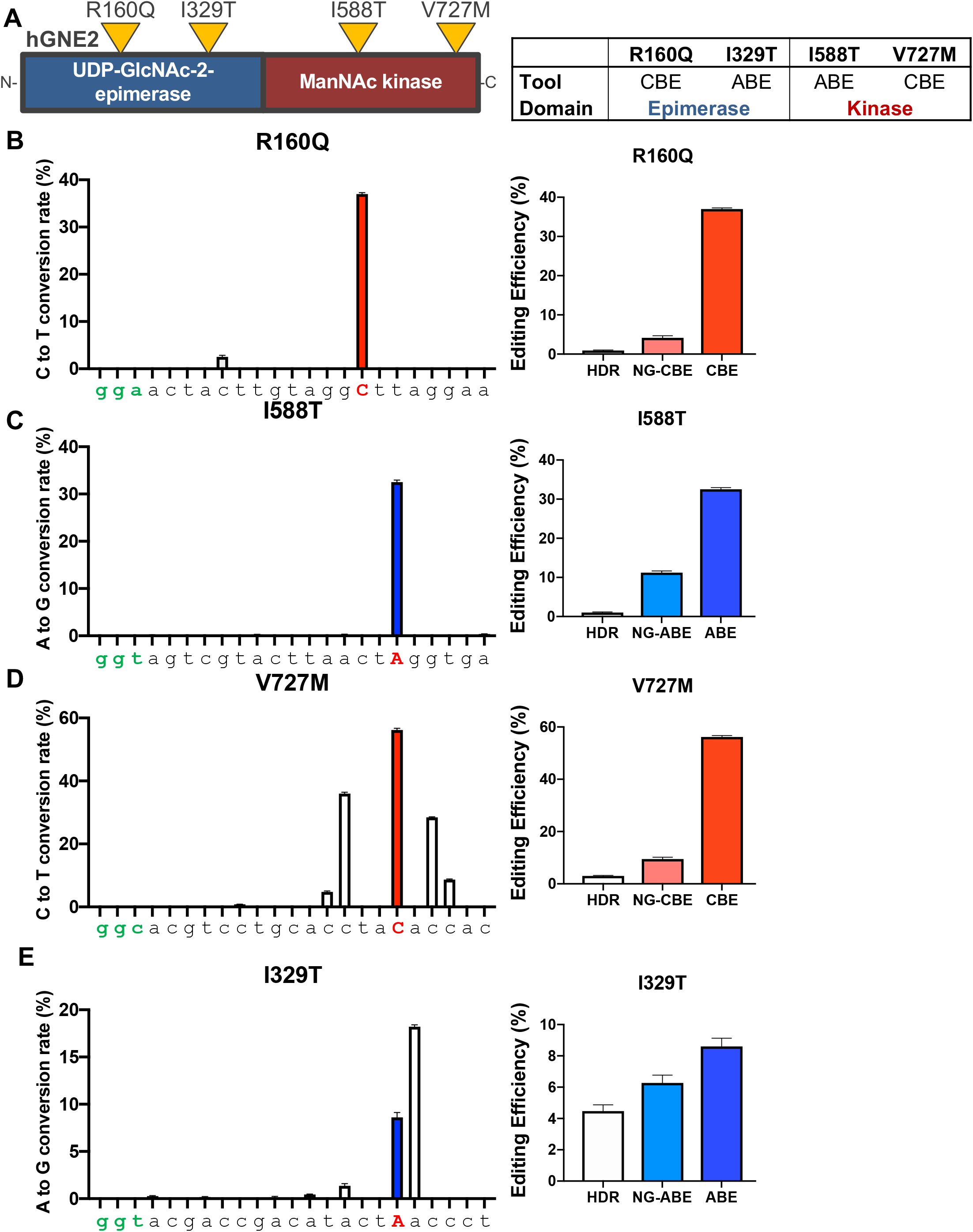
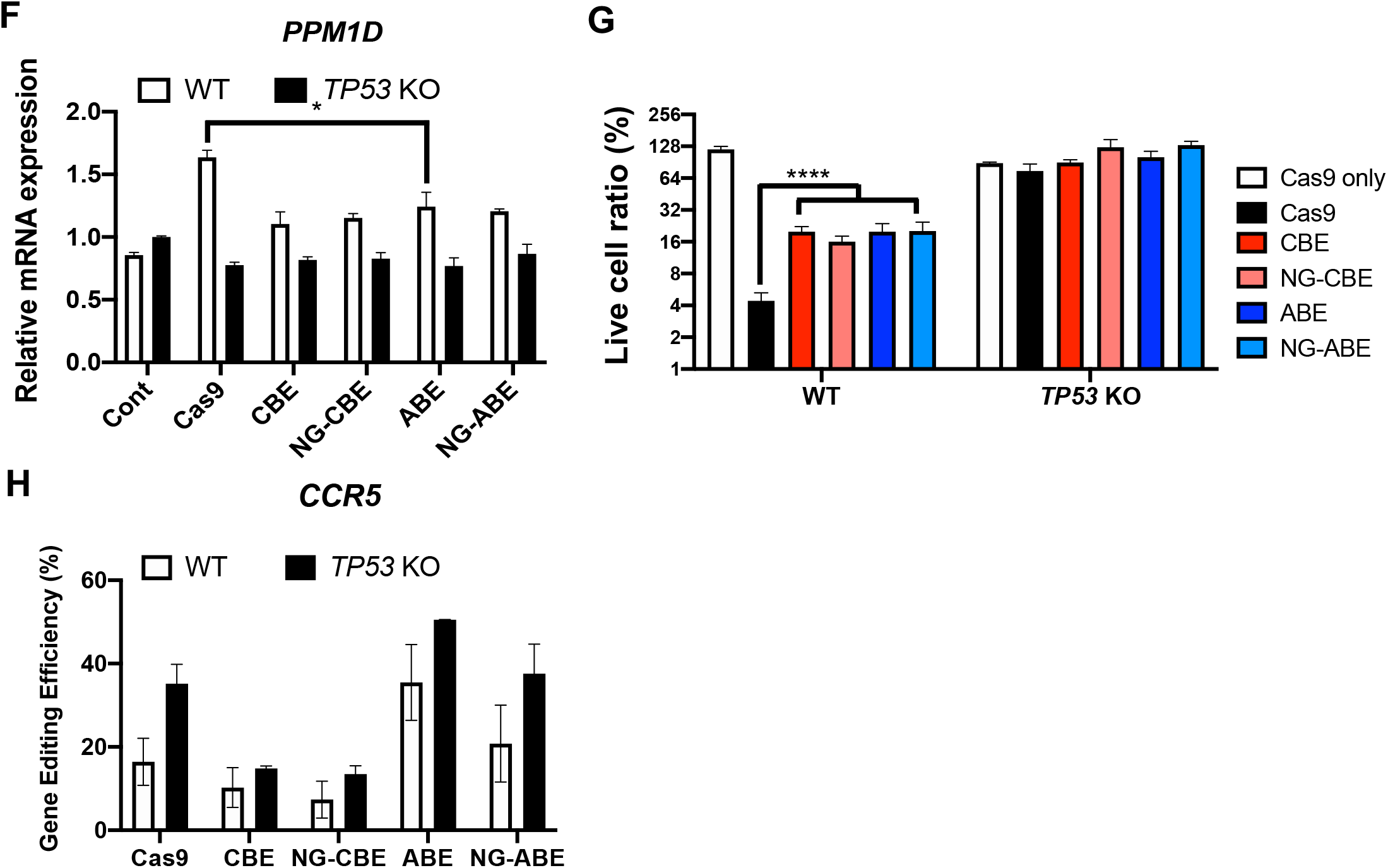
Efficiency of base editing compared to HDR in hESCs. (A) Graphical description of site of mutations in the epimerase and kinase domain of human GNE protein (hGNE2) (left), List of mutations generated by each base editor (right) (B to E) Target deep sequencing results showing overall base conversion rate of each sequence by CBE (C to T) and ABE (A to G) (left), Graphical presentation of on-target editing efficiency of indicated gene editing tools (right) for R160Q (B), I588T (C), V727M (D) and I329T (E), Target base and PAM sequence is colored in red and green (3’-5’) respectively. (F) mRNA expression of *PPM1D* in wild-type (WT) and *TP53* Knockout (TP53 KO) hESCs transfected with sgRNA targeting CCR5 (sgCCR5) and indicated gene editing tools (Cas9, CBE, NG-CBE, ABE and NG-ABE) compared to control (Cont: sgCCR5 only), *, p<0.05 (G) Graphical presentation of percentage of live cell ratio of wild-type (WT) and TP53 KO hESCs after transfection of sgCCR5 and indicated gene editing tools, ****, p<0.0001 (H) Gene editing efficiency of indicated gene editing tools in WT and TP53 KO hESCs on CCR5. Editing efficiencies are analyzed by target deep sequencing.

As low HDR efficiency compared to indel by Cas9 for GNE mutations (Fig. S1G) demands laborious clonal selection in hPSCs (Li et al., 2016), the inefficiency of HDR remains a serious technical limitation to be resolved for disease modeling in hPSCs despite the advantage of broader target-ability. Furthermore, we observed that NGG-BEs (ABE and CBE) were more efficient than NG-BEs (NG-ABE and NG-CBE) as previously reported (Jeong et al., 2019).

### P53 response by Cas9 determines editing efficiency

The extremely low gene editing efficiency with Cas9 in hPSCs (Kim et al., 2020; Mali et al., 2013) results from p53-dependent cell death upon DSB caused by the endonuclease activity of Cas9 (Ihry et al., 2018). hPSCs are particularly susceptible to p53-dependent cell death once DSB occurs, which is thought to be a protective mechanism to rigorously maintain genome integrity during development (Weissbein et al., 2014). Thus, we hypothesized that the low editing efficiency of HDR compared to BEs (Fig. 2) may result from different p53 responses. To explore this hypothesis, we first established *TP53* KO hESCs by targeting exon 3 of *TP53* (Fig. S2A). After introducing sgRNA and Cas9 in hESCs, Nutlin3, which is an inhibitor of MDM2, was applied to stabilize p53, causing massive cell death, as expected (Fig. S2B). After Nutlin3 treatment, the few clones that survived were *TP53* KO hESCs (Fig. S2C). The p53-dependent response of Cas9 or BEs was assessed by the measuring the level of *PPM1D*, a typical target gene of p53 (Fiscella et al., 1997). When introducing Cas9 with sgRNA into hESCs as previously described (Ihry et al., 2018), BEs only marginally affected the p53 response (Fig. 2F). Consistent with the p53 response, significantly more hESCs remained and survived when BEs were used, compared to the Cas9 approach, while all *TP53* KO hESCs tolerated the stress of gene editing (Fig. 2G). The effects of gene editing techniques in WT and *TP53* KO hESCs were monitored in real-time (Movies S1 and S2). Along with the higher rate of survival in *TP53* KO hESCs than in wild-type hESCs (Fig. 2G), improvement of the gene editing efficiency of Cas9 was evident in *TP53* KO hESCs (Fig. 2H).

### Establishment of GNE myopathy disease models through base editors

Even with the limited target-ability of BEs (Figs. 1C and D), significant higher gene editing efficiency of BEs than HDR may enable BEs to be currently the most feasible approach for base substitution in hPSCs. Accordingly, we successfully established 14 GNE mutants in total in both hESCs and hiPSCs using BEs (Table S2). Briefly, based on H9 hESCs, two LOF mutants in the epimerase domain of GNE generated with CBE (R160Q) and ABE (I329T) (Figs. 3A and B) and two LOF mutants in the kinase domain using NG-ABE (I588T) and CBE (V727M) were established (Figs. 3C and D). Other GNE mutant cell lines (I329T, R160Q, and I588T) were established in CHA3-hESCs (Fig. S3D) and BJ-iPSCs (Figs. S3E and F) using NG-ABE, ABE, or CBE, as listed in Table S2. Of note, since the target windows of I329T and V727M carry bystander bases, five mutants with bystander mutations including I329I, a silent mutation, were also established in V727M (Figs. S3B and C) and I329T (Fig. S3D).

**Figure 3.**
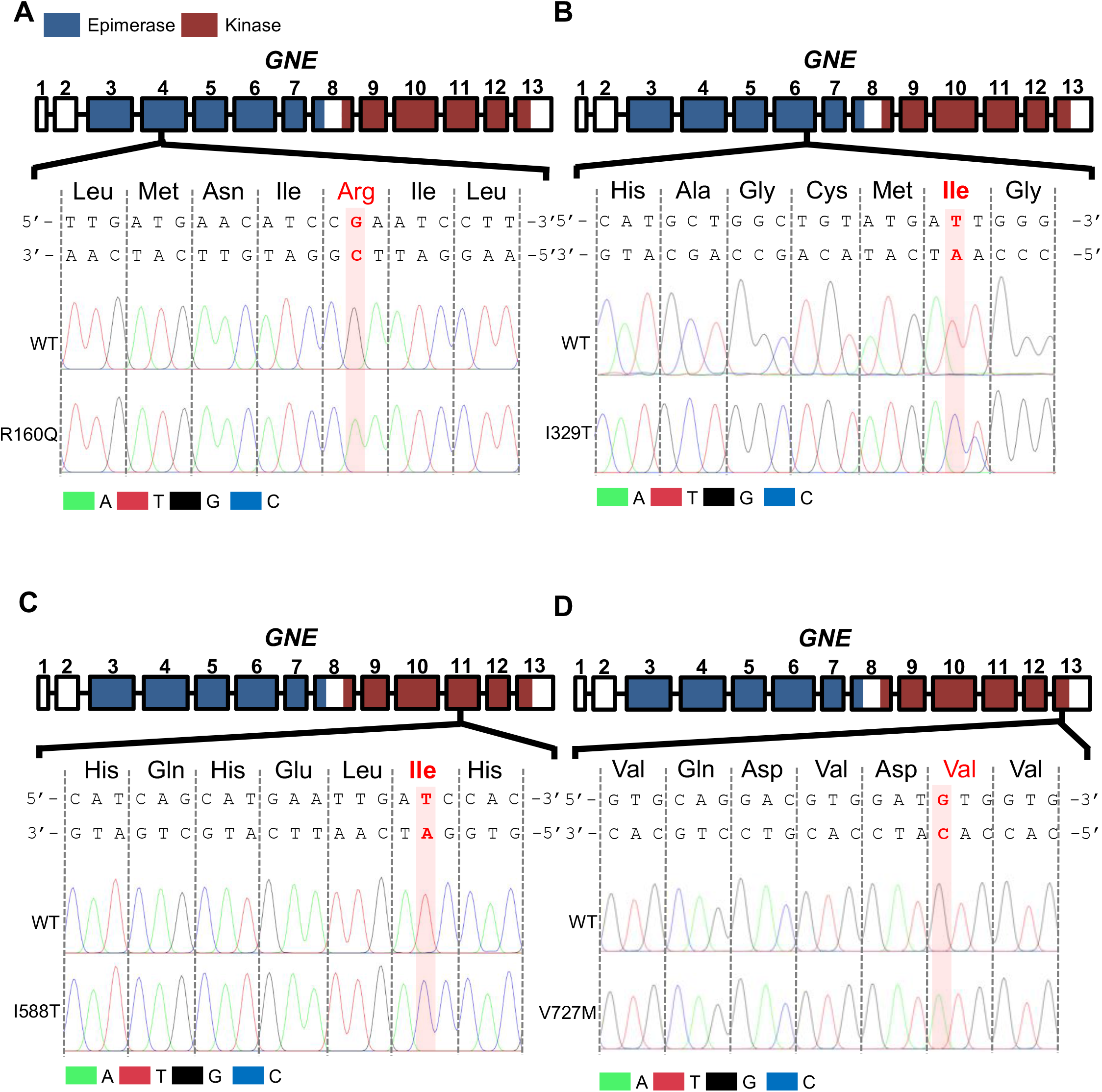
Establishment of GNE myopathy disease models through base editors. (A to D) Sanger validation of mutants of R160Q (A), I329T (B), I588T (C) and V727M (D) compared to wild-type (WT), Exons coding UDP-GlcNAc-2-epimerase and ManNAc kinase domain are colored in blue and in red respectively. Target amino acids and bases are colored in red.

### Mutation-specific phenotype of GNE myopathy disease models

The four established GNE mutant hESCs (GNE-hESCs) displayed typical cellular morphology and comparable OCT4 expression (Fig. 4A). As the putative underlying cause of GNE myopathy is hyposialylation, we first assessed levels of sialic acid production using a sialic acid-specific *Sambucus nigra agglutinin* (SNA) lectin binding assay in four different GNE-hESCs. As shown in Figure 4B, the levels of lectin staining were significantly diminished in all mutant hESCs to different degrees, relative to controls. The mean fluorescence intensity (MFI) of SNA lectin clearly indicated that the sialylation of the protein varied depending on the mutation (Figs. 4C and D). Of note, binding of SNA lectin indicates the level of sialylation of protein, the final step for sialic acid biosynthesis, is a reliable indicator of GNE enzymatic activity. In this line, MFI of SNA staining in GNE mutants relative to WT (Fig. 4D) can be used to determine pathogenicity. Thus, we compared previously predicted severity of each mutations by *in silico* platforms (PolyPhen2, SIFT, Align, and Pmut) (Celeste et al., 2014) to the MFI of each GNE mutant (Fig. 4E). I329T mutant, predicted as “severe” revealed the most obvious hyposialylation (Figs. 4D and E). As the V727M mutation occurs in the compound heterozygous state, even with the prevalent distribution of this variant (rs121908627), this mutation itself would have a less serious effect on GNE myopathy according to the moderate hyposialylation in the V727M mutant (Fig. 4C). We would expect the I588T mutant, which is a variant of uncertain significance (VUS), to be “Pathogenic/Likely pathogenic” since hyposialylation levels in this mutant are comparable to those observed in the R160Q mutant, which is classified as “Pathogenic/Likely pathogenic” (Fig. 4E). The hyposialylation of other GNE models derived from BJ-iPSC and hCHA3 hESCs were examined. Evident hyposialylation (Fig. S4A) and recovery by sialic acid treatment (Fig. S4B), compared to that of the silent I329I mutation established in hCHA3 cells, ruled out the possibility of baseline effects of gene editing itself on hyposialylation. The comparable degree of hyposialylation of R160Q and I588T was also reproduced in GNE mutants derived from BJ-iPSCs (Figs. S4C and D). Importantly, the levels of sialic acid in four different hPSC lines (two hESCs and two iPSCs), differed in a cell type-specific manner depending on genetic background (Fig. 4F) despite comparable level of mRNA expression of *GNE* (Fig. 4G). This supports the necessity of an isogenic pair for precise comparison as highlighted previously (Merkle and Eggan, 2013).

**Figure 4.**
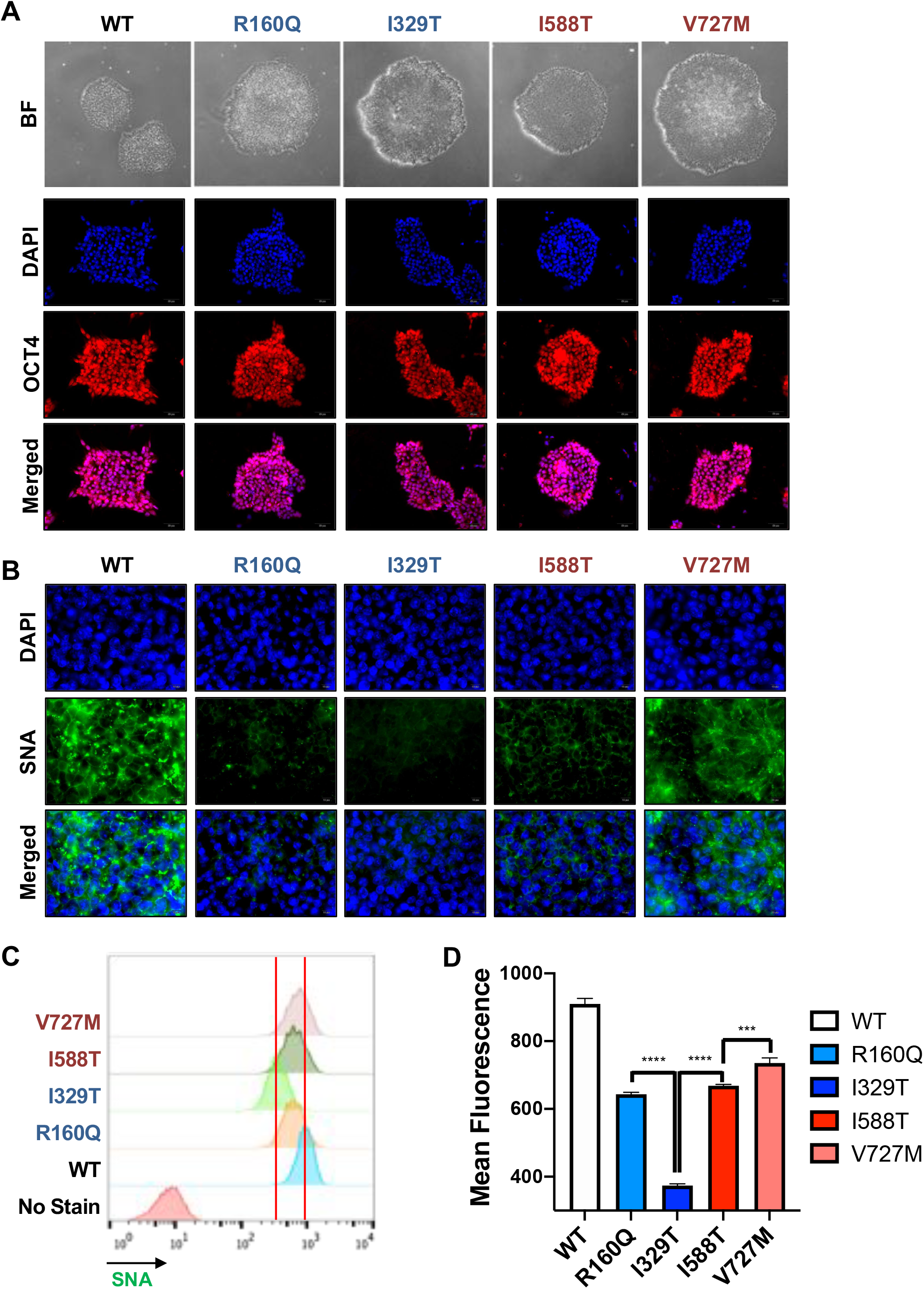
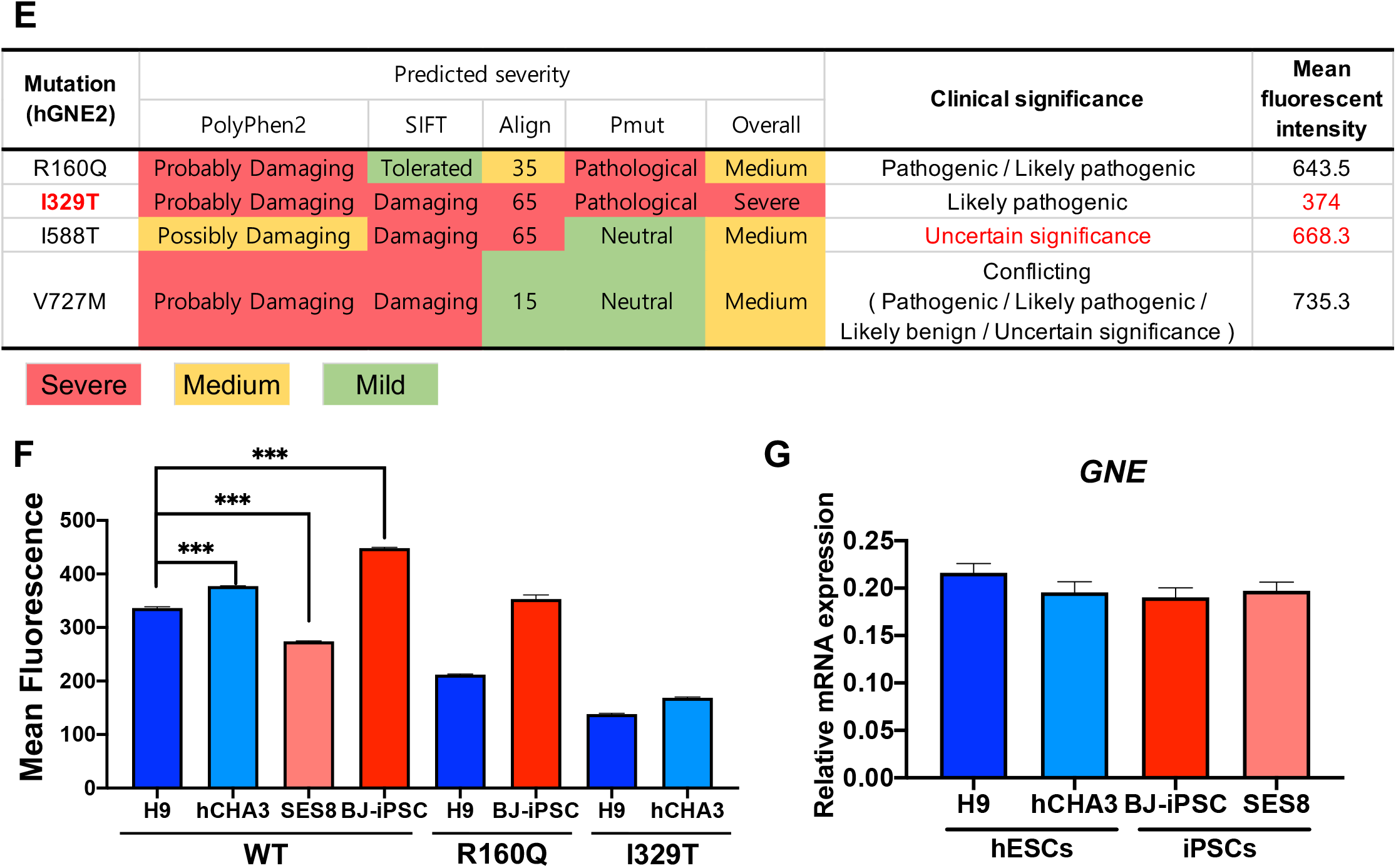
Mutation-specific phenotype of GNE myopathy disease models. (A) Microscope bright field (BF) images (top) and fluorescent images for OCT4 (bottom) of hESCs of WT and GNE mutants, DAPI for nuclear counterstaining (B) Fluorescent images of hESCs of WT and GNE mutants stained with SNA (green) and DAPI for counter nuclear staining (C) Histogram and (D) bar graph of mean fluorescence intensity (MFI) of SNA in WT and GNE mutants, ***, p<0.001, ****, p<0.0001 (E) List of predicted severity, clinical significance and mean fluorescence intensity of GNE mutants, predicted severe, medium and mild were colored in red, yellow and green respectively. (F) Mean fluorescence intensity of SNA of indicated hPSCs lines, ***, p < 0.001 (G) mRNA expression of *GNE* in indicated hPSCs lines

### Structural analysis predicts mutation-specific hyposialylation

The varying degrees of hyposialylation among the GNE mutants inspired us to analyze each mutation at the molecular level. We used the GNE structure to predict the molecular consequences of each mutation on the enzymatic activity of GNE. The epimerization reaction of GNE requires tetramerization through the N-terminal domain, which is stabilized upon ligand binding (Fig. 5A) (Chen et al., 2016; Ghaderi et al., 2007). R160 stabilizes the C-terminal loop via interaction with the main chains of D409, Q411, and E412 (Fig. 5B). This loop interacts with the residues involved in the N-terminal dimerization interface. The R160Q mutant contains a shorter side chain that would require loop reorganization were they to interact, which could possibly affect epimerase oligomerization (Fig. 5B). However, the hyposialylation effect of R160Q was less adverse than that of the I329T mutation. I329 participates in forming an extensive network of hydrophobic interactions (Fig 5C). Importantly, L249, located adjacent to I329, interacts directly with uridine diphosphate (UDP) (Fig. 5C). The I329T mutation, which introduces a smaller and less hydrophobic residue, would weaken the Van der Waals interactions within the hydrophobic core, affecting the structural integrity near the substrate-binding site. Meanwhile, the kinase domain of GNE displays a typical bi-lobal architecture and contains a type I zinc-binding motif characteristic of the repressor, open reading frame, kinase (ROK) family (Fig. 5A) (Martinez et al., 2012; Tong et al., 2009). Full-length GNE requires dimerization to permit ManNAc kinase activity (Martinez et al., 2012). I588 is located at the dimerization interface of the kinase C-terminal lobe (Fig. 5D); mutating this residue to less-hydrophobic threonine (Thr) affects kinase dimerization and thus its activity. Finally, V727 is buried in a hydrophobic core (Tong et al., 2009). The modeling of the V727M mutation suggests that the bulkier methionine (Met) may induce structural changes at this site to avoid steric clash (Fig. 5E). However, since this residue is positioned away from the catalytic site, the V727M mutation is expected to cause a less severe phenotype than the I329T mutation.

**Figure 5.**
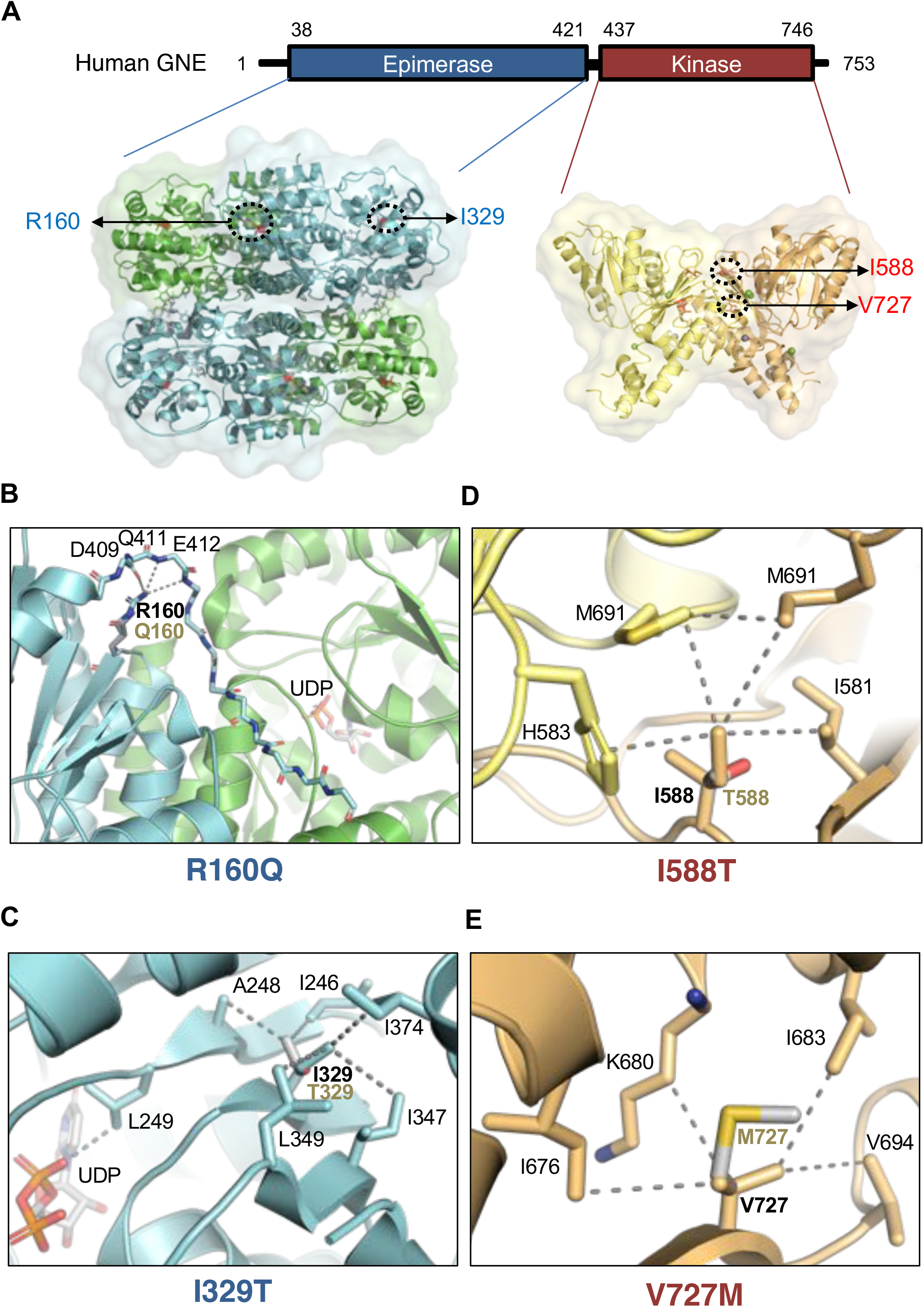
Structural analysis predicts mutation-specific hyposialylation. (A) Structures of tetrameric epimerase and dimeric kinase domains of human GNE (PDB ID 4ZHT and 2YHW, respectively) are shown. Target mutations R160Q (B) and I329T (C) are found in the epimerase domain, while I588T (D) and V727M (E) are in the kinase domain. Mutagenesis and rotamer selection were performed using PyMOL, and mutated residues are indicated in grey. Inter-residue interactions within a distance of 4.1 Å are indicated with dashed lines.

Structural analysis of GNE mutants shows that the I329T mutation, which can cause structural instability, is located near the active site. As a result, this mutation is expected to have the most adverse effect on GNE epimerase activity. Indeed, the I329T mutant shows the most pronounced hyposialylation (Figs. 4C and D).

### Mutation-specific drug response of GNE myopathy disease models

To compensate for the hyposialylation caused by reduced GNE enzymatic activity in GNE myopathy patients, administration of ManNAc is being investigated in clinical trials for GNE patients (NCT02346461) following a safety profile study (Xu et al., 2017), despite the unsatisfactory outcome of a sialic acid clinical trial (Lochmuller et al., 2019). We therefore hypothesized that ManNAc would elicit a distinctive response in GNE patients with a mutation in the domain of ManNAc kinase because ManNAc is a direct substrate of this enzyme (Fig. 6A). To explore this possibility, we examined the recovery of hyposialylation of GNE mutants in the epimerase domain (R160Q and I329T) and kinase domain (I588T and V727M) after treatment with ManNAc. As predicted, the recovery of sialic acid production after ManNAc treatment occurred in GNE mutants harboring mutations in the epimerase domains (R160Q and I329T), while the effect in the kinase mutant (I588T and V727M) was marginal (Figs. 6B and C). The dose-dependent response curve clearly revealed the mutant-specific ManNAc response (Figs. 6D and E). We noticed that sialic acid production was reduced in V727M, for unknown reasons (Fig. 6E). These data suggest that the ManNAc response differs according to the mutation type of GNE.

**Figure 6.**
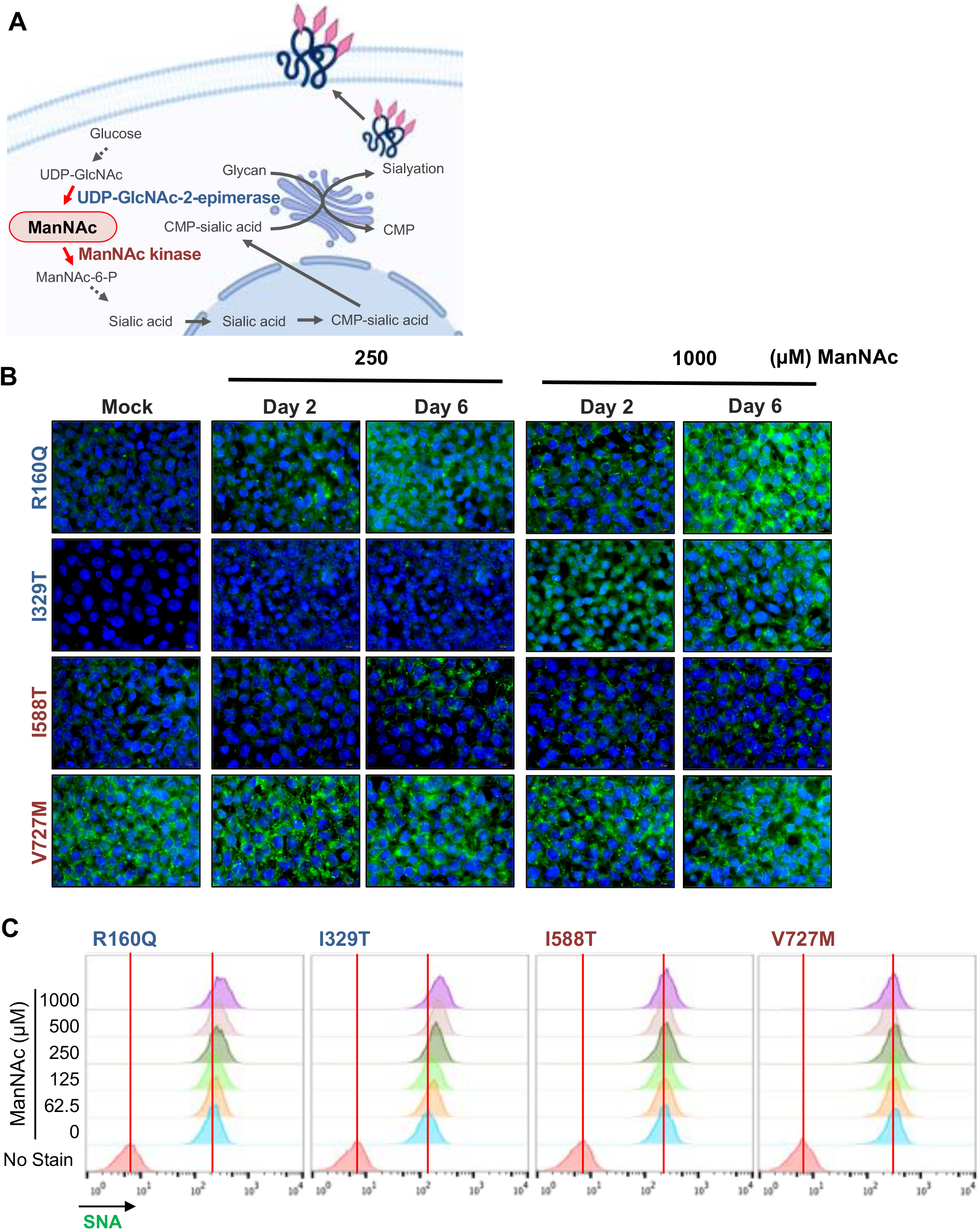
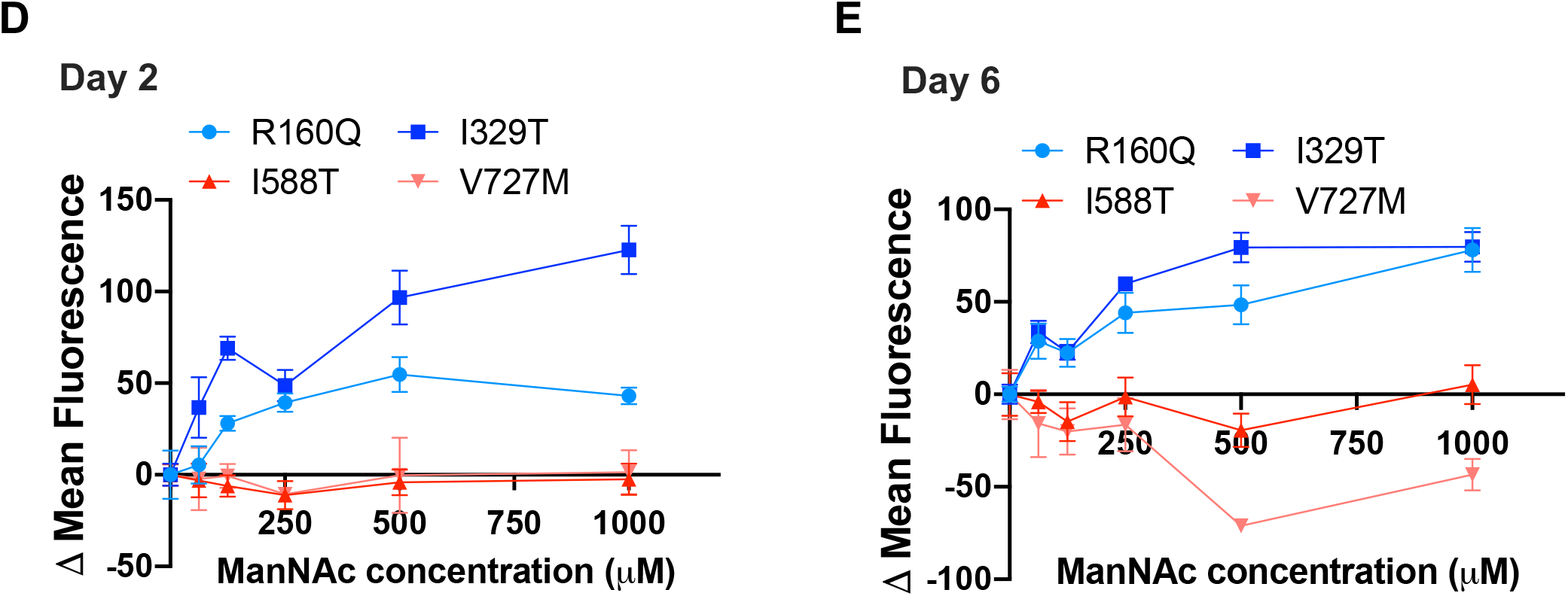
Mutation-specific drug response of GNE myopathy disease models. (A) Graphical description of biosynthesis of sialic acid and sialyation and roles of GNE protein as UDP-GlcNAc-2-epimerase and ManNAc kinase (B) Fluorescence microscopic images of GNE mutant hESCs stained with SNA after indicated dose of ManNAc treatment for day 2 and day 6 (C) Flow cytometry for SNA in GNE mutant hESCs 6 days after indicated dose of ManNAc treatment (D and E) Change of mean fluorescent intensity of SNA at 2 (D) and 6 (E) days after indicated concentration of ManNAc treatment

### GNE myopathy modeling in myocytes derived from GNE-hESCs

Prior to inducing the differentiation of GNE-hESCs into myocytes, we examined whether hyposialylation would affect “pluripotency.” To examine this effect, I329T hESCs were used, as they showed the most severe hyposialylation. Neither alkaline phosphatase activity, which is a well-characterized pluripotency indicator, nor the mRNA expression of typical pluripotency markers were altered (Figs. S5A and B). Additionally, teratomas from the I329T mutant developed normally (Fig. S5C). Thus, we concluded that hyposialylation due to GNE mutation had minimal effects on the pluripotency of hESCs. Next, to verify mutation-specific responses in myocytes, WT and GNE-hESCs were differentiated into skeletal myocytes (hereafter referred to as myocytes) through multiple steps (Fig. 7A). The expression levels of typical markers for myocytes (e.g., *MYOG, MYH2*, and *MYH7*) and pluripotency (e.g., *NANOG* and *SOX2*) were used to evaluate myocyte differentiation (Fig. 7B). Differentiated myocytes showing typical elongated, rod-shaped morphology, clearly expressed MYH3 (also known as Myosin heavy chain-embryonic: MyHC-emb), a skeletal muscle specific contractile protein (Fig. 7C). Notably, the mutation-specific hyposialylation observed in GNE-hESCs (Fig. 4D) was also evident in GNE-myocytes relative to controls (Figs. 7D and E). Consistently, exclusive effect of ManNAc on myocytes with epimerase mutation (R160Q and I329T) was also reproduced (Figs. 7F and G). While understanding of the functionality of skeletal myocytes derived from hPSCs is limited, the functionality of cardiomyocytes differentiated from hESCs (hESC-CMs) has been well-characterized (Mummery et al., 2012). Thus, we examined whether GNE mutation affects the functionality of cardiomyocytes, as cardiomyopathy has been occasionally reported in GNE patients (Malicdan et al., 2014). hESC-CMs with higher Troponin T2, Cardiac type (*TNNT2*) and lower *NANOG* expression than hESCs (Fig. S6A) showed clear beating behavior (Movie S3). GNE mRNA expression was comparable regardless of mutation status (Fig. 6B). Consistent mutation-specific hyposialylation and drug responses were also observed in hESCs-CMs with different GNE mutations (Figs. S6C). Consistently, mutation specific ManNAc response was reproduced in hESC-CMs (Fig. S6D). The hyposialylation of hESC-CMs with the I329T mutation was evident when compared with hESCs-CM with the I329I silent mutation (Fig. S6E). The comparison of beating behavior of hESC-CM from WT or I329T hESCs revealed no significant changes resulting from GNE mutation (Movies S4A and B).

**Figure 7.**
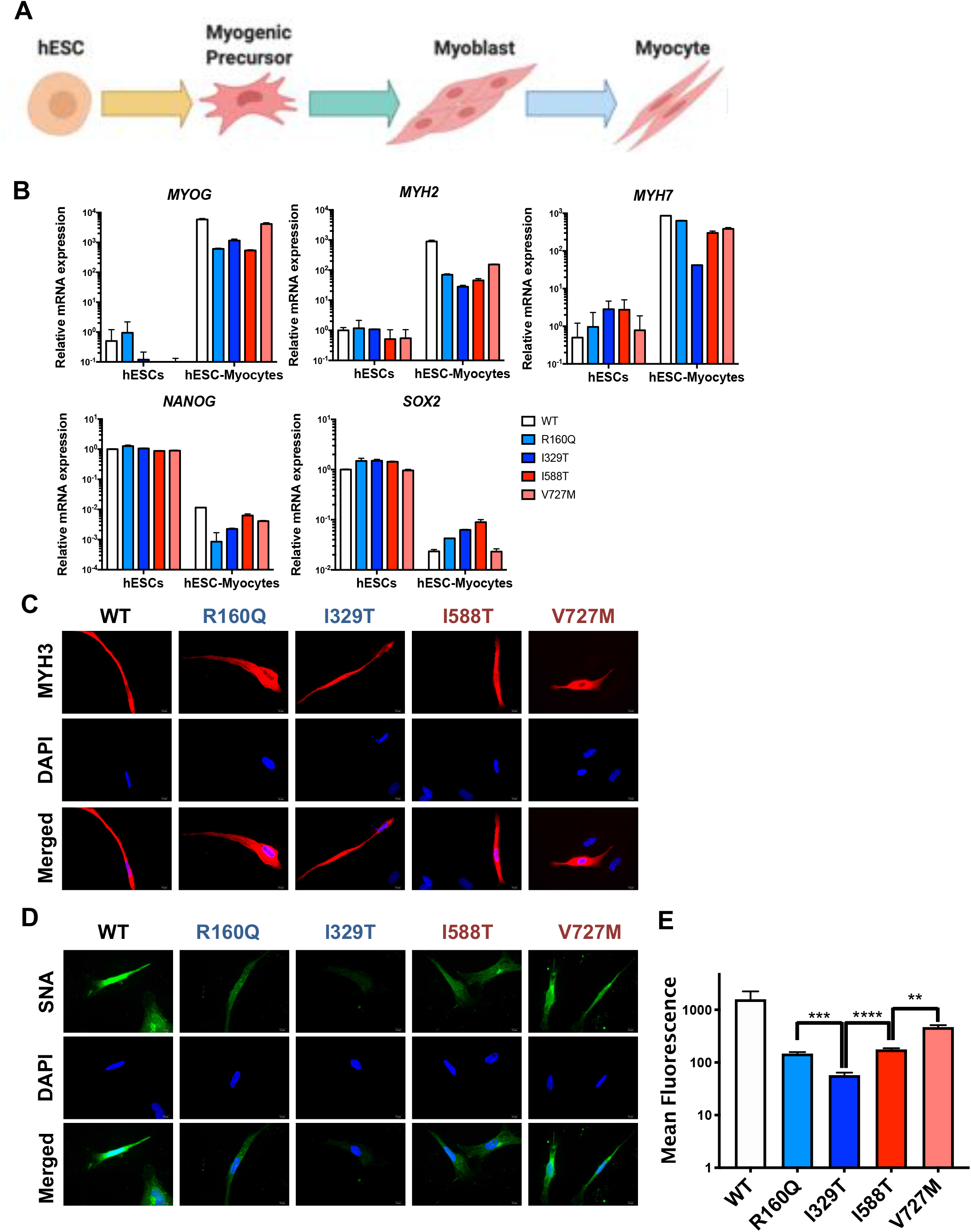
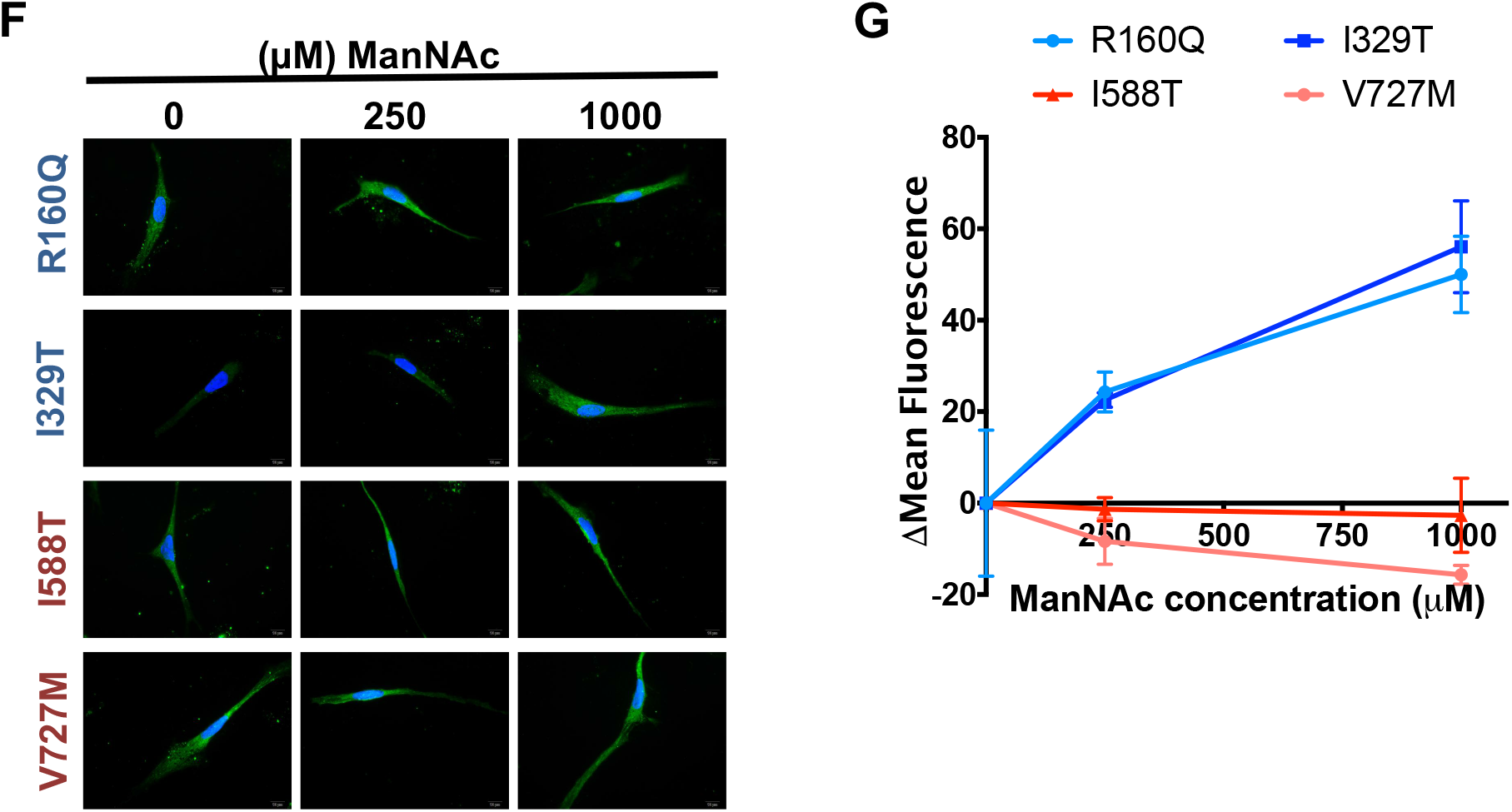
GNE myopathy modeling in myocytes derived from GNE-hESCs. (A) Schematic diagram of myogenic differentiation from hESCs (B) mRNA expression of typical markers for skeletal myocytes (top) and pluripotency (bottom) in hESCs and myocytes derived from hESCs of WT and GNE mutants (C) Fluorescence microscope images for MYH3 in myocytes derived from WT and GNE mutant hESCs, DAPI for nuclear counterstaining (D) Fluorescent images of hESCs derived myocytes of WT and GNE mutants stained with SNA (green) and DAPI for counter nuclear staining (E) bar graph of mean fluorescence intensity (MFI) of SNA in WT and GNE mutants, **, p<0.01, ***, p<0.001, ****, p<0.0001 (F) Fluorescence microscopic images of GNE mutant hESCs derived myocytes stained with SNA after indicated dose of ManNAc treatment (G) Change of mean fluorescent intensity of SNA after indicated concentration of ManNAc treatment

## Discussion

### Base editors for disease modeling in hPSCs

Despite the limited target accessibility of BEs relative to HDR, the high editing efficiency of BEs compared to HDR in hPSCs would be advantageous in disease modeling using hPSCs. Through the development and implementation of NG-BEs for gene editing, coverage of BEs for GNE myopathy modeling encompassed 73.5% of transition point mutations in the present study (Fig. 1). We demonstrated that BEs were more efficient than HDR due to their lesser effect on p53 (Figs. 2F and H). However, along with PAM-less target sequences, which account for 26.5% of transition mutations in GNE myopathy, the production of undesirable mutations of bystander bases with BEs, would be a significant drawback in disease modeling. Thus, the development of “PAM-less BEs” and “BEs with narrowed editing windows” would be advantageous for more efficient disease modeling with BE techniques.

### Significance of isogenic disease model

Here, we established 14 GNE mutants from three hPSCs with different genetic backgrounds. Hyposialylation, a typical pathological phenotype of GNE myopathy, occurred in a mutation-specific manner regardless of hPSC cell type. However, because the basal level of sialic acid in the wild-type of the four hPSCs (H9, hCHA3: hESCs, BJ-iPSCs and SES8 (Lee et al., 2010): iPSCs) differed relative to the mutant hPSCs (Fig. 4F), comparison of sialic acid between unmatched pairs could not allow us to precisely determine the effect of mutation. Previously, various disease models were established using iPSCs derived from patients, in which disease phenotype was compared to genetically unmatched control iPSCs (Lee et al., 2009; Yamashita et al., 2014). This study confirmed the previously proposed bias, introduced by the use of cells with an unmatched genetic background to hinder precise comparisons of pathological phenotypes [4].

### Base editing technique for validating a variant of uncertain significance

Despite large numbers of single nucleotide variants (SNVs) in rare genetic diseases, the functional consequences of mutations are often unpredictable, due to insufficient *a priori* observations and incomplete penetrance (Findlay et al., 2018). With isogenic disease models, one can classify these so-called “variants of uncertain significance” (VUS) (Findlay et al., 2018) more accurately with the aid of functional information. In the present study, for example, since I588T showed a similar level of hyposialylation as R160Q, classified as “Pathogenic/Likely pathogenic,” we inferred that I588T may also be pathogenic. The I588T variant was originally annotated as “uncertain significance” in the ClinVar database (Landrum et al., 2016). The efficient induction of point mutation with BEs (or PE in the near future) would enhance further studies seeking to validate the effect of VUS.

### Consideration of mutation-specific drug response

We also demonstrated that epimerase mutants (R160Q and I329T) but not kinase mutants (I588T and V727M) showed recovery of sialylation after ManNAc treatment (Figs. 6D and 7G). Interestingly, while the I588T mutant showed no recovery, the V727M mutation resulted in reduced sialylation following ManNAc treatment, for unknown reasons. In light of these mutant-specific responses to ManNAc, it would be advisable to use patient stratification based on mutation status in clinical trials of ManNAc, to ensure accurate efficacy validation with ManNAc and similar drugs.

### Future prospects for drug development based on isogenic disease models

A recent study developed a high throughput (HT) phenotypic screening system to measure CFTR protein in cystic fibrosis with an isogenic pair of iPSCs (Merkert et al., 2019). Similarly, along with 3D protein modeling (Fig. 5), HT phenotypic screening with isogenic GNE myopathy disease models using the level of sialylation as a phenotypic indicator could accelerate drug development for GNE myopathy. Furthermore, by taking advantage of isogenic disease models and a drug-transcriptome database such as the Connectivity MAP (Kwon et al., 2019), candidate drugs from the FDA-approved drug library that reverse pathogenic gene signatures can easily be predicted (Kwon et al., 2020; Liu et al., 2015). The efficacy of these drug candidates can then be validated in the “GNE myopathy in a dish” model, making this strategy a strong potential option to develop drugs for GNE myopathy in the future.

### Conclusion

The isogenic “GNE myopathy in a dish model,” established in hPSCs by BEs with relatively high efficiency, revealed mutation-specific hyposialylation and drug responses. This work establishes important cellular tools and techniques for further mechanistic studies and development of drug screening systems.

## Supporting information

Supplementary informations

## Author contributions

HJ.C conceived the overall study design and led the experiments. JC.P and J.K mainly conducted the experiments, data analysis, and critical discussion of the results. HK.J and S.B contributed to sgRNA design and NGS analysis and provided gene editing techniques. SY.L, KT.K and S.P. produced mutant lines. HS.L and HJ.C performed 3D protein structure modeling. SJ.P and SH.M conducted the experiments with differentiated cardiomyocytes.

### Declaration of competing interest

The authors declare that they have no known competing financial interests or personal relationships that could have appeared to influence the work reported in this paper.

## Acknowledgments

This work was supported by a grant from the National Research Foundation of Korea (2017M3A9B3061843 and 2020R1A2C2005914) and a grant from the Research of Korea Centers for Disease Control and Prevention (2020-ER6902-00).

## Materials and Methods

### Live cell ratio after gene editing in WT and TP53 KO hESCs

To trace the number of live cells during gene editing, cells were documented by JuLI stage for 36 hours after transfection. Cells were transfected with the same condition in ‘Generation of *TP53* knock-out and *GNE* mutant hPSCs’. The number of cells at the time point of 2 hours and 36 hours were counted by cell counter ImageJ plugin. To calculate the live cell ratio, the number of cells at the time point of 36 hours were divided by the number of cells at the time point of 2 hours.

### Generation of TP53 knock-out and GNE mutant hPSCs

For hPSCs transfection, cells were detached with Accutase™ (561527, BD Bioscience) and washed with DPBS for three times and resuspended to concentration of 1 × 10^6^ cell in 100 μl with Opti-MEM (31985070, Gibco). 2 μg of Cas9 or BE vectors (Cas9, BE4max, NG-BE4max, ABEmax and NG-ABEmax cloned in pCMV vector), 3 μg of sgRNA vector and 2 μg of ssODN (in case of HDR) was added to 100 μl of cell mixture. Electroporation was performed by NEPA-21 with 175 V of poring pulse and 2.5 mV of transfer pulse. To enrich *TP53* KO cells, 10 μM of nutlin3 was treated for 48 hours after 5 days from transfection.

### Flow cytometry

Cells were detached with Accutase solution (561527, BD Bio-sciences) for hPSCs or with TrypLE (12604, Gibco) for myocytes, followed by 3 times DPBS wash and then analyzed through FACS Calibur I (BD Bioscience). For the determination of sialic acid, cells were stained with Fluorescein SNA (FL-1301, Vector Laboratories). CellQuest Pro soft-ware was used for FACS analysis.

### Generation of skeletal muscle cells from hESCs

hESCs were differentiated to skeletal muscle cells by using commercially available skeletal muscle differentiation medium kit (Amsbio). In brief, Cells were seeded onto a collagen I (Sigma)-coated dish at 5,000 cells /cm2 dish and maintained for 6 days in skeletal muscle induction medium (SKM01, Amsbio). At day 6, cells were dissociated with 0.05% trypsin, seeded onto a collagen I-coated dish at 5,000 cells /cm2 dish, and maintained for 6 days in myoblast cell culture medium (SKM02, Amsbio). At day 12, when myoblasts are confluent, the medium was changed to myotube cell culture medium (SKM03, Amsbio) and freshly changed every 3 days for more than 6 days. For drug response test, cells were treated with ManNAc (Sigma) at the indicated times and concentrations and the media with ManNAc was changed every day.

### Statistical analysis

The quantitative data are expressed as the mean values ± standard deviation (SD). Student’s unpaired t-tests was performed to analyze the statistical significance of each response variable using the PRISM. p values less than 0.05 were considered statistically significant (*, p < 0.05, **, p <0.01, ***, p <0.001 and ****, p <0.0001)

## Graphical abstract

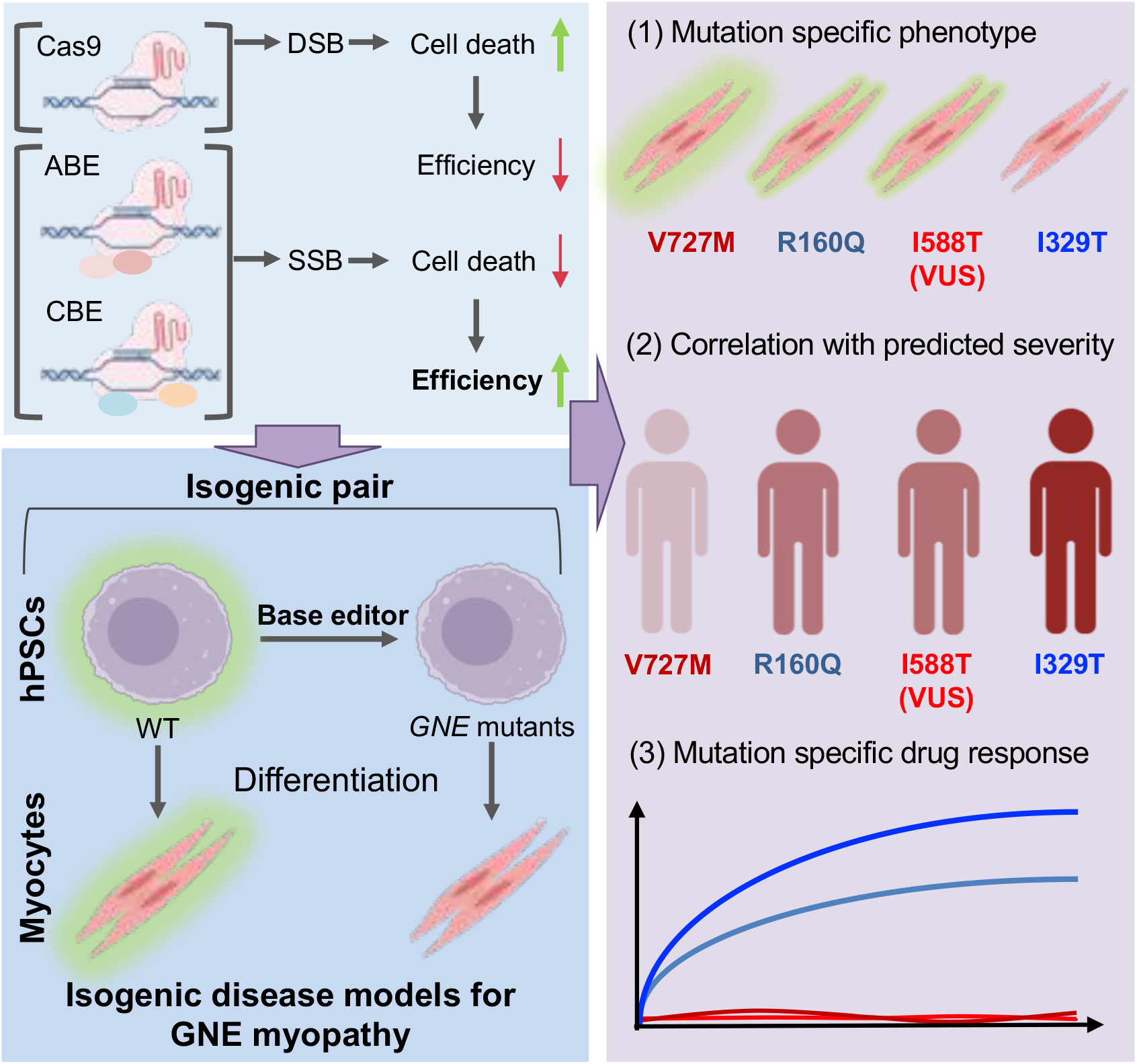

## Notes

### Competing Interest Statement

The authors have declared no competing interest.

