## Supplementary informations for "Efficient GNE myopathy disease modeling with mutation specific phenotypes in human pluripotent stem cells by base editors"

This PDF file includes:

Experimental Section

Supplemental Figure legends

Supplementary Figures 1-6

#### Supplementary Materials and Methods

##### ClinVar

An excel file including variants associated with GNE myopathy was downloaded from ClinVar database (https://www.ncbi.nlm.nih.gov/clinvar/). Total 63 mutations that annotated in ClinVar as "pathogenic" or "likely pathogenic" were selected, and classified into mutation types (e.g., transition point mutation, transversion point mutation, indel, duplication or others). Among 34 transition point mutations, whether the mutations can be a target for base editors is determined by checking if they have Protospacer Adjacent Motif (PAM) sequence for base editors.

##### Cell line and culture

Human embryonic stem cells (WA09, WiCell Research Institute and hCHA3) and induced pluripotent stem cells (SES8 and BJ-iPSCs) were cultured on Matrigel (Corning, #354277) coated dishes fed with mTeSR-E8 media (STEMCELL technologies) and StemMACS media (Miltenyi-Biotec) added with 50 μg/ml Gentamicin (Gibco). Cells were passaged every 5–6 days and media was changed every day. hPSC were washed with DPBS and detached with Dispase (Gibco). Detached cells were rinsed with DMEM/F-12 media (Gibco) and plated on Matrigel-coated dish. 10 μM of Y27632 (Gibco) was added for cell attachment when needed. For drug response test, cells were treated with ManNAc (Sigma) at the indicated times and concentrations and the media with ManNAc was changed every day.

##### Teratoma formation assay

As described previously [49], $5x10^6$ cells of each WT and GNE I329T hESC were injected into the testes of 4-weeks-old male BALB/C nude mice (n =3). Mice were euthanized 10 weeks after injection. These animal experiments were conducted under the permission of Seoul National University Institutional Animal Care and Use Committee (Permission number: SNU-180810-1-1).

**Immunocytochemistry**

Cells were treated with cold methanol for fixation and permeabilization and treated with 3% BSA in PBS for blocking. For lectin staining, Fluorescein SNA (FL-1301, Vector Laboratories) and DAPI (Sigma-Aldrich, #D9542) was diluted in blocking solution and applied to cells for 1 hour at room temperature, dark condition. Fluorescein SNA and DAPI were washed off three times with PBS and mounted on slide glass using MOWIOL solution. Fluorescence microscopy (Olympus) was used for imaging samples.

**RT-qPCR analysis**

Easy-BLUE™ RNA isolation kit (iNtRON Biotechnology) is used for total RNA extraction. PrimeScript™ RT reagent kit (TaKaRa) is used to generate cDNA from RNA extracted previously. Quantitative real-time PCR analysis was performed with Light Cycler-480®II (Loche) and SYBR® Green PCR reagents (Life Technologies), following the supplier's instructions.

**Generation of cardiomyocytes from hESC**

Myocyte differentiation from hESCs was performed as previously described [50]. In brief, cells were seeded onto a hPSC qualified Matrigel (Corning, #354277)-coated cell culture dish at 140,000 cells/cm2 dish. 10μM Y-27632 (Gibco) was added for the first 24 hours after passage. The medium was changed daily, and cells were allowed to grow in StemMACS for 3–4 days until the cells were 90% confluent. At day 0, cells were treated with 4 μM CHIR99021 (#2520691, Peprotech) in DM1 [Differentiation medium 1: RPMI1640 (ThermoFisher Scientific)/Insulin-free B27 Supplement (Gibco)]. After 24 hours, the medium was changed to DM1 supplemented with 3 μM CHIR99021 for additional 24 hours. At day 2, the medium was changed with 2 μM C59 (Wnt inhibitor, #1248913, Peprotech) in DM1. After 24 hours, the medium was replaced with 1 μM C59 in DM1. At day 4, the medium was changed with DM1. At day 6, the medium was changed with DM2 [Differentiation medium 2: Advanced MEM (Gibco)/GlutaMax (Gibco)] andn freshly changed every 2 days for more than 4 days. For drug response test, cells were treated with ManNAc (Sigma) at the indicated times and concentrations and the media with ManNAc was changed every day.

**Alkaline Phosphatase Staining**

Alkaline phosphatase (AP) staining was performed with Alkaline Phosphatase Kit (86R1KT, Sigma Diagnostics™), following the supplier's instructions.

**Targeted deep-sequencing analysis**

Genomic DNA from each samples was isolated using Wizard® genomic DNA purification kit (Promega). To assess gene editing, targeted deep sequencing near the DNA cleavage site was performed. Briefly, genomic DNA was prepared from the electroporated rotifer population, and the targeted regions (~ 300 bp) were amplified by adaptor primers with Phusion polymerase (New England Biolabs, Radnor, PA, USA). The detailed protocol was described in a previous study (Park et al., 2017). The resulting amplicons were subjected to paired-end read sequencing using MiniSeq (Illumina, San Diego, CA, USA). After MiniSeq, paired-end reads were joined by the Fastq-join utility, a part of the ea-utils program (https://code.google.com/archive/p/ea-utils/), using default values. Paired-end reads were then analyzed by comparing wild type and mutant sequences using CasAnalyzer (Park et al., 2017).

**Sequence information**

**1. PCR primer information**

| Primer pairs for sanger sequencing | Primer sequence (5' - 3') |
| --- | --- |

| | | |
|---|---|---|
| **GNE_R160Q** | **F1** | GAGAATCGCTTGAACCCGG |
| | **F2** | TTGCATGTTACGGCTCTGTG |
| | **R1** | ACAGGGCAAGGTCTATGAACT |
| | **R2** | AACCTTGGCCTTCTCATTGC |
| **GNE_I329T** | **F1** | GGTTCGAGTGATGCGGAAGA |
| | **F2** | ATGCGGAAGAAGGGCATTGA |
| | **R1** | AACCTCATTCACTCACTTGTTCAC |
| | **R2** | GCCTAGTCAGTTGAAGTATGAGC |
| **GNE_I588T** | **F1** | CCATCCAAAATGCTGCCAGAG |
| | **F2** | GCTTTGATGCCTAGTGGGCT |
| | **R1** | AGTTTTAGTGCGAGGGAGCC |
| | **R2** | CCATTCCAGAGGCGTATGCT |
| **GNE_V727M** | **F1** | GGTACACCCGGCTTTTTCCA |
| | **F2** | GAACAGCTTTGGGTCTTGGG |
| | **R1** | TCTCTGCCAAAGTCACCTGC |
| | **R2** | AGGTCCATGTCTGTTCCTGG |

**2. qPCR primer information**

| **Primer Pairs for qPCR** | | **Primer Sequence (5'→3')** |
|---|---|---|
| 18s rRNA | F | GTAACCCGTTGAACCCCATT |
| | R | CCATCCAATCGGTAGTAGCG |
| PPM1D | F | CAGACACAAGTGTCCACACT |
| | R | TGGCACAAGATTGTCCATGC |
| GNE | F | GCAGGGAGCAAAGAGATGGT |
| | R | TCTGACGTGTTCCCAGGTTG |
| SOX2 | F | TTCACATGTCCCAGCACTACCAGA |
| | R | TCACATGTGTGAGAGGGGCAGTGTGC |
| LIN28 | F | CACGGTGCGGGCATCTG |
| | R | CCTTCCATGTGCAGCTTACTC |
| MyoG | F | GGGCGTGTAAGGTGTGTAAGA |
| | R | AGGTTGTGGGCATCTGTGTAGG |
| MYH2 | F | GAAATTACTTCTGGCAAAATACAGG |
| | R | GCAGATGCCAGTTTTCCAGT |
| MYH7 | F | CACCAACAACCCCTACGATT |
| | R | ACTCATTGCCCACTTTCACC |
| NANOG | F | AAATTGGTGATGAAGATGTATTCG |
| | R | GCAAAACAGAGCCAAAAACG |

| | | |
|---|---|---|
| TNNT2 | F | ATGAGCGGGAGAAGGAGCGGCAGAAC |
| | R | TCAATGGCCAGCACCTTCCTCCTCTC |

**3. sgRNA sequence information**

| Gene | gRNA sequence (with NGG PAM) |
|---|---|
| **CCR5** | **GGCAGCATAGTGAGCCCAGAAGG** |
| **TP53** | **CCCTTCCCAGAAAACCTACCAGG** |
| **GNE_R160Q** | **AAGGATTCGGATGTTCATCAAGG** |
| **GNE_I329T** | **TCCCAATCATACAGCCAGCATGG** |
| **GNE_I588T** | **CGTGGATCAATTCATGCTGATGG** |
| **GNE_V727M** | **CACCACATCCACGTCCTGCACGG** |

**4. Donor DNA sequence information**

| Gene | ssODN sequence |
|---|---|
| **R160Q** | **TTA ATC GCC TGA AGC CTG ATA TCA TGA TTG TTC ATG GAG ACA GGT TTG ATG CCC TGG CTC TGG CCA CAT CTG CTG CCT TGA TGA ACA TCC AAA TCC TTC ACA TTG AAG GTG GGG AAG TCA GTG GGA C** |
| **I588T** | **ATA GTT TTC TGT TTC AGG TAA AGC ACC CTC AGT GAT TTC TAT CCC TCG CTG TAG GAA TCG GTG GTG GAA TTA TCC ATC AGC ATG AAT TGA CCC ACG GAA GCT CCT TCT GTG CTG CAG AAC TGG GCC A** |
| **I329T** | **AGG GCA TTG AGC ATC ATC CCA ACT TTC GTG CAG TTA AAC ACG TCC CAT TTG ACC AGT TTA TAC AGT TGG TTG CCC ATG CTG GCT GTA TGA CTG GGA ACA GCA GCT GTG GGG TTC GAG AAG TTG GAG C** |
| **V727M** | **ATC CTC TCC GGA GTC CTG GCC AGT CAC TAT ATC CAC ATT GTC AAA GAC GTC ATT CGC CAG CAG GCC TTG TCC TCC GTG CAG GAC GTG GAT ATG GTG GTT TCG GAT TTG GTT GAC CCC GCC CTG CTG G** |

Antibody information

1. Primary antibody

| Protein | Company | Catalog # |
|---|---|---|
| OCT4 | Abcam | 19857 |
| MYH3 | DSHB | F1.652 |

2. Secondary antibody

| Product name | Company | Catalog # |
|---|---|---|

| | | |
|---|---|---|
| anti-mouse, Alexa Fluor 594 | Invitrogen | A-11005 |

3. Lectin for FACS

| Protein | Company | Catalog # |
|---|---|---|
| SNA-FITC | Vector Laboratories | FL-1301 |

**Supplemental Figure legends**

**Figure S1.** (A to D) Overall base conversion rate of each sequence for R160Q (A), I588T (B), V727M (C), I329T (D), Target base and PAM sequence is colored in red and green (3'-5') respectively. (E and F) Sequence of each genotype produced in (E) V727 and (F) I329 site and frequency by CBE and NG-CBE, Target base and bystander bases are colored in red and in green respectively. Pathogenic mutations are colored in red. (G) Graphical presentation of indel (black) and HDR (red) frequency in each indicated targets

**Figure S2.** (A) Graphical presentation of site (exon 3) of *TP53* targeting gRNA (gTP53), proline-rich domain and DNA binding domain are colored in red and blue respectively. (B) Bright field microscopic images of hESCs after introducing gTP53 and Cas9 with 10 μM of Nutlin3 treatment, *TP53* KO hESCs remained survived are indicated by red arrow. (C) Indel size spectrum (x-axis) and frequency (y-axis) of *TP53*, achieved by next generation sequencing (left), List of insertion and deletion of wild-type (WT) and pool of *TP53* KO hESCs (right)

**Figure S3.** (A to D) Sanger sequencing data of R160R/Q (A), V727I, D726D/N and V727V/M (B) , V727M/I, V727M / D726D/N and V727M / D726N (C) from H9 hESCs, (D) Sanger sequencing data of I329I and I329T from hCHA3 hESCs (E and F) Sanger sequencing data of R160Q (E) and I588T (F) from BJ-iPSCs

**Figure S4.** (A) Fluorescent images I329I (top) and I329T (bottom) hESCs stained with SNA, DAPI for nuclear counterstaining (left), Flow cytometry of SNA staining in I329I and I329T hESCs (right) (B) Fluorescent images of I329T hESCs with vehicle (Mock) or sialic acid (SA, 1mM), DAPI for nuclear counterstaining (left), Flow cytometry of SNA staining in I329I and I329T hESCs at 48 hours after treatment of 1mM of sialic acid (SA) (C) Fluorescent images of wild-type (WT), R160Q and I588T mutants from

BJ-iPSCs stained with SNA, DAPI for nuclear counterstaining (D) Graphical presentation of relative mean fluorescence intensity of SNA in wild-type (WT), R160Q and I588T mutants from BJ-iPSCs

**Figure S5.** (A) Microscope images of wild-type (WT) and I329T hESCs after alkaline phosphatase activity assay (left) and images of colonies after alkaline phosphatase activity assay (right) (B) Relative mRNA expression of typical pluripotent markers in wild-type (WT) and I329T hESCs (C) Microscopic images H&E staining of typical three germ layers in the teratoma from I329T hESCs, The scale bars represent 200μm (top) and 50 μm (bottom).

**Figure S6.** (A and B) Relative mRNA expression of cardiomyocyte markers (A) and *GNE* (B) in WT hESC and cardiomyocytes (CMs) derived from GNE mutant hESCs. (C) Fluorescent images of CMs of WT and GNE mutants stained with SNA, DAPI for nuclear counterstaining (D) Change of mean fluorescent intensity of SNA after indicated concentration of ManNAc treatment in R160Q, I329T and I588T mutant hESCs derived CMs 48 hours after indicative concentration of ManNAc treatment. (E) Fluorescent images of CMs derived from I329I and I329T stained with SNA, DAPI for nuclear counterstaining (left), Flow cytometry of SNA of CMs derived from I329I and I329T (right)

**Movie S1.** Real-time bright filed image of WT hESCs transfected with designated editing tools with gRNA targeting CCR5

**Movie S2.** Real-time bright filed image of TP53 KO hESCs transfected with designated editing tools with gRNA targeting CCR5

**Movie S3.** Beating cardiomyocytes derived from wild-type hESC.

**Movie S4.** (A) Beating cardiomyocytes of WT at day 14 of differentiation (B) Beating cardiomyocytes of GNE I329T at day 14 of differentiation

**Table S1.** List of nucleotide and protein change, clinical significance from ClinVar database and available editing tool for each mutation

**Table S2**. List of GNE mutant cell lines established by designated base editing tools

**Figure. S1**

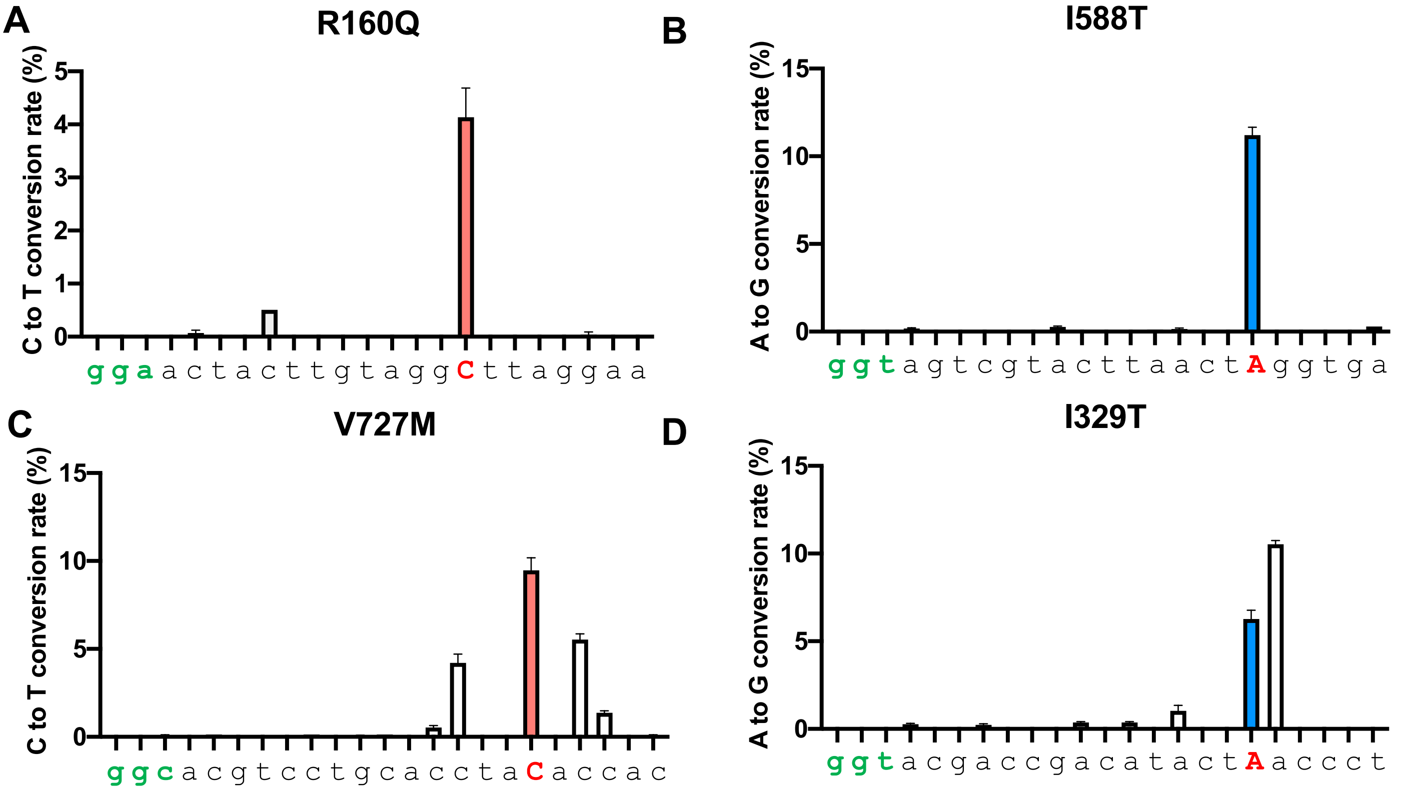


**E**

| gRNA sequence | Genotype | Frequency (%) CBE | Frequency (%) NG-CBE |
|---|---|---|---|
| C A C C A C A T C C A C G T C C T G C A C G G | WT | 32.54 | 88.21 |
| C A C C A T A T C C A C G T C C T G C A C G G | **V727M** | 13.45 | 3.54 |
| C A C C A T A T T C A C G T C C T G C A C G G | V727M; D726N | 14.40 | 1.22 |
| C A C T A T A T C C A C G T C C T G C A C G G | V727I | 9.18 | 2.04 |
| C A C C A C A T T C A C G T C C T G C A C G G | D726N | 4.74 | 0.83 |
| C A T T A T A T C C A C G T C C T G C A C G G | V727I; V728M | 3.30 | 0.49 |
| C A T T A T A T T C A C G T C C T G C A C G G | D726N; V727I; V728M | 1.76 | 0.31 |
| C A C T A C A T C C A C G T C C T G C A C G G | V727V | 1.66 | 0.70 |

**F**

| gRNA sequence | Genotype | Frequency (%) CBE | Frequency (%) NG-CBE |
|---|---|---|---|
| T C C C A A T C A T A C A G C C A G C A T G G | WT | 74.70 | 84.32 |
| T C C C A G T C A T A C A G C C A G C A T G G | **I329T** | 6.79 | 4.90 |
| T C C C G G T C A T A C A G C C A G C A T G G | **I329T** | 1.45 | 1.11 |
| T C C C G A T C A T A C A G C C A G C A T G G | I329I | 15.58 | 8.84 |

**G**

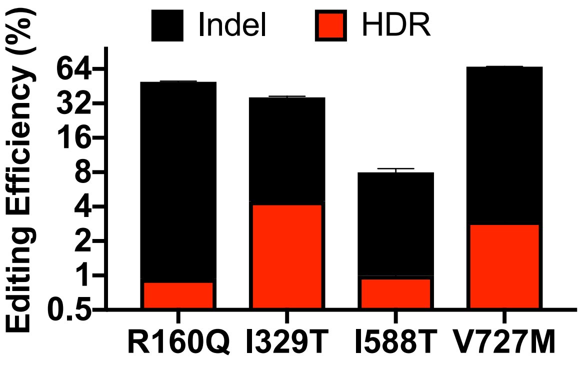

### Figure. S2

**A**

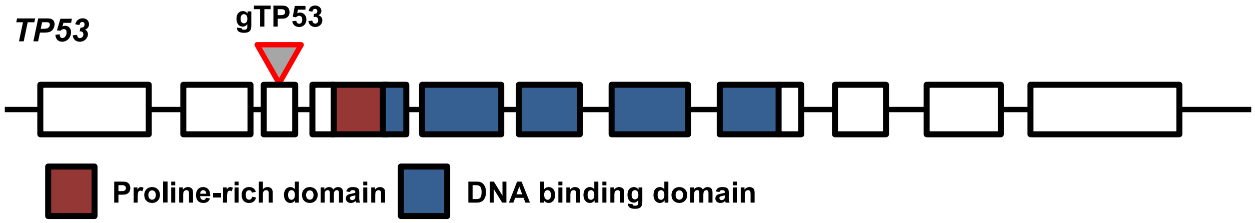


**B**

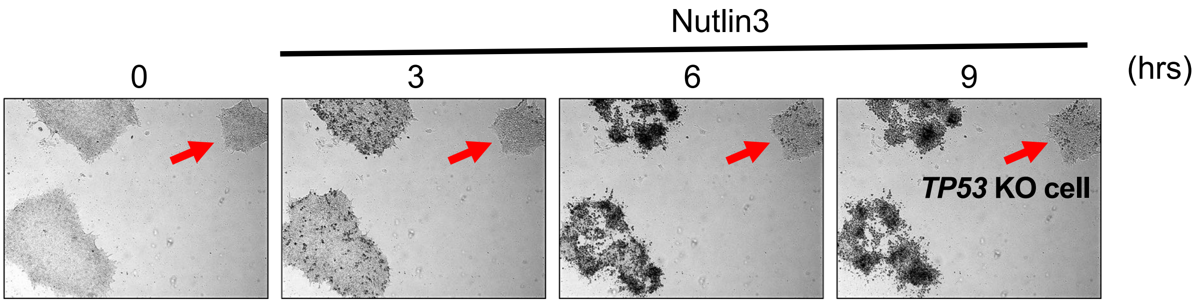


**C**

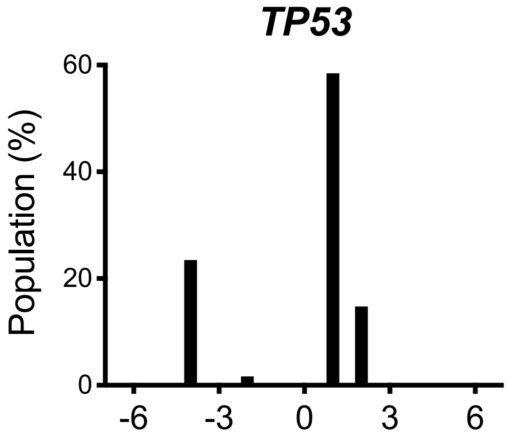


| Index | Group | Reads | Ins | Del | Indel |
|---|---|---|---|---|---|
| 32 | WT | 6599 | 11 | 7 | 18 (0.3%) |
| 33 | KO (Pool) | 6692 | 4741 | 1944 | 6685 (99.9%) |

Figure. S3

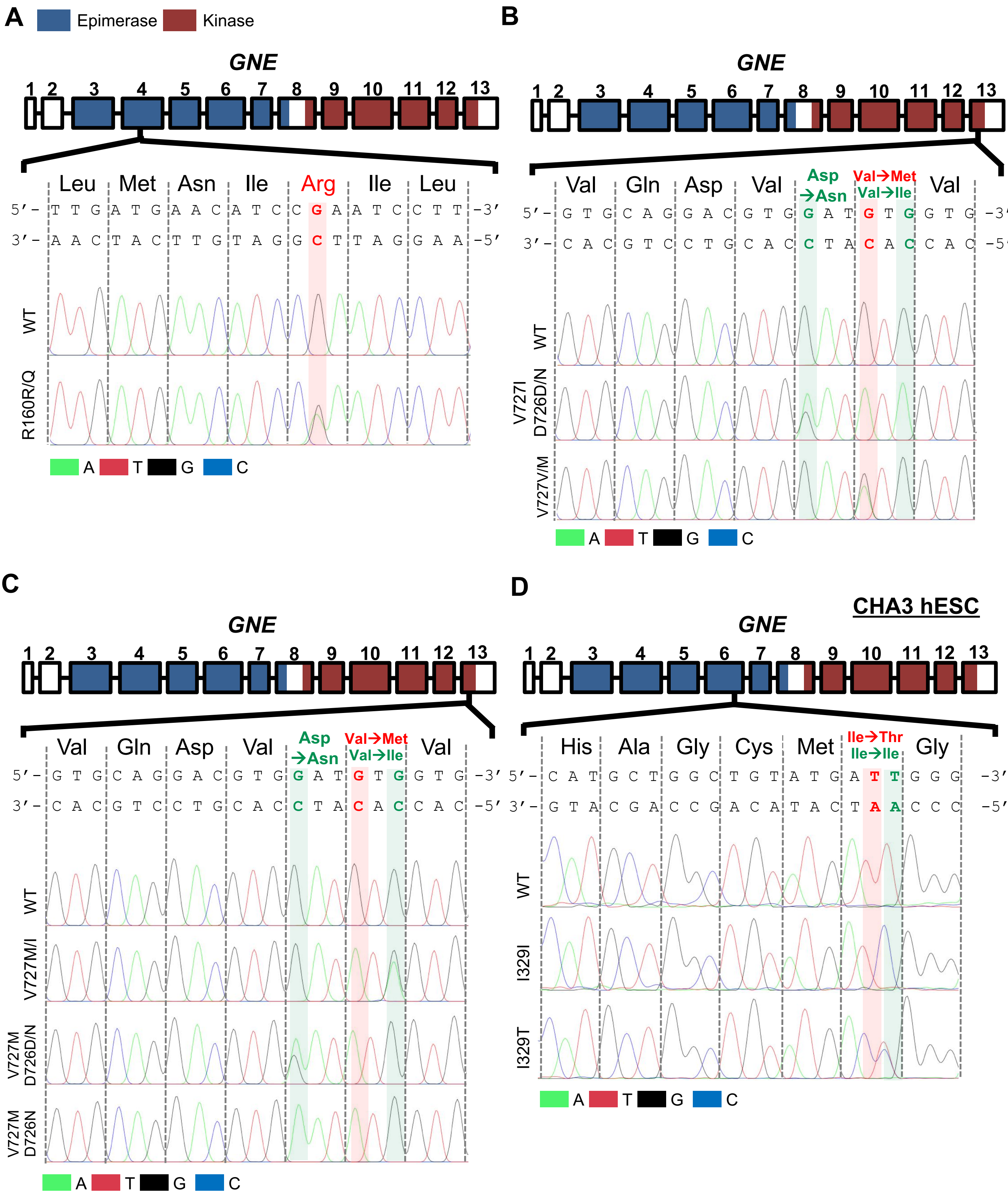
A
Epimerase
Kinase
GNE
1 2 3 4 5 6 7 8 9 10 11 12 13
Leu Met Asn Ile Arg Ile Leu
5′- T T G A T G A A C A T C C G A A T C C T T -3′
3′- A A C T A C T T G T A G G C T T A G G A A -5′
WT
R160R/Q
A T G C
B
GNE
Val Gln Asp Val Asp →Asn Val→Met Val→Ile Val
5′- G T G C A G G A C G T G G A T G T G G T G -3′
3′- C A C G T C C T G C A C C T A C A C C A C -5′
WT
V727I D726D/N
V727V/M
A T G C
C
GNE
Val Gln Asp Val Asp →Asn Val→Met Val→Ile Val
5′- G T G C A G G A C G T G G A T G T G G T G -3′
3′- C A C G T C C T G C A C C T A C A C C A C -5′
WT
V727M/I
V727M D726D/N
V727M D726N
A T G C
D
CHA3 hESC
GNE
His Ala Gly Cys Met Ile→Thr Ile→Ile Gly
5′- C A T G C T G G C T G T A T G A T T G G G -3′
3′- G T A C G A C C G A C A T A C T A A C C C -5′
WT
I329I
I329T
A T G C

Figure. S3

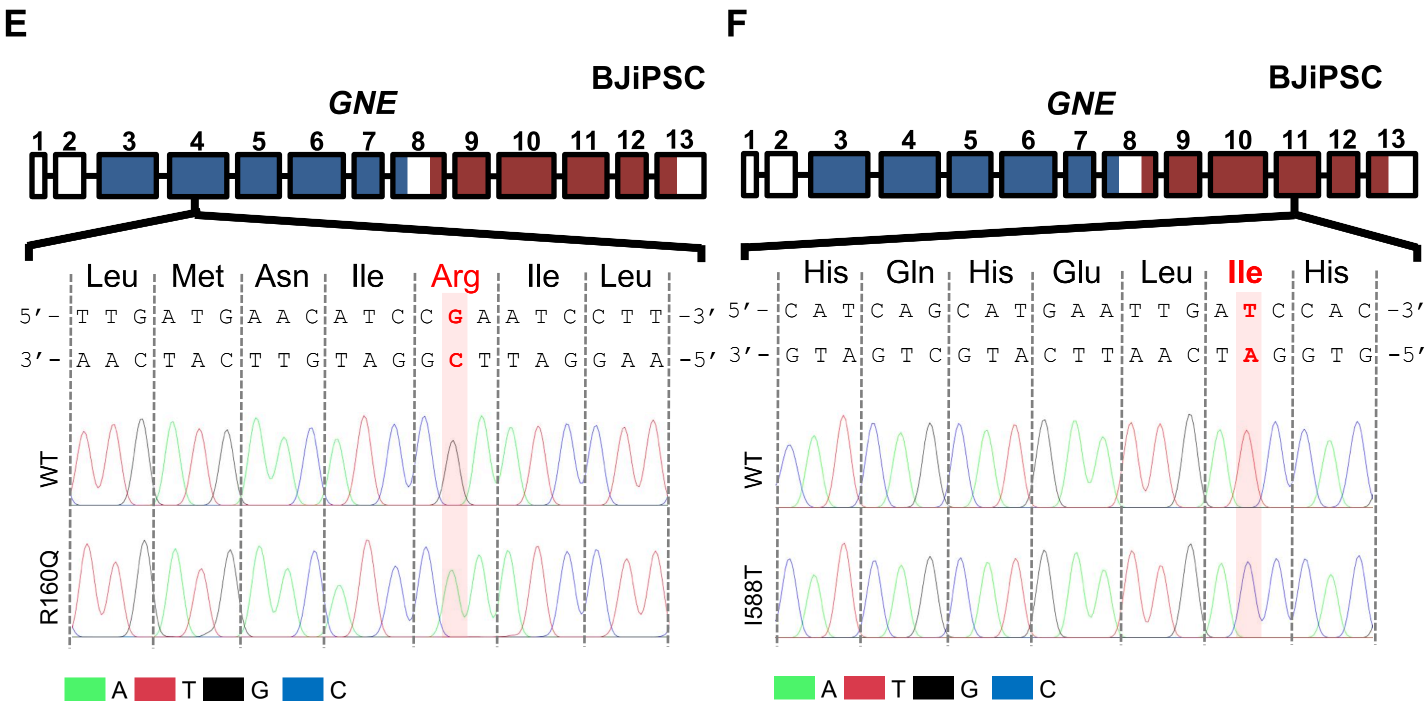

E
BJiPSC
GNE
1 2 3 4 5 6 7 8 9 10 11 12 13
Leu Met Asn Ile Arg Ile Leu
5′- T T G A T G A A C A T C C G A A T C C T T -3′
3′- A A C T A C T T G T A G G C T T A G G A A -5′
WT
R160Q
A T G C
F
BJiPSC
GNE
1 2 3 4 5 6 7 8 9 10 11 12 13
His Gln His Glu Leu Ile His
5′- C A T C A G C A T G A A T T G A T C C A C -3′
3′- G T A G T C G T A C T T A A C T A G G T G -5′
WT
I588T
A T G C

Figure. S4

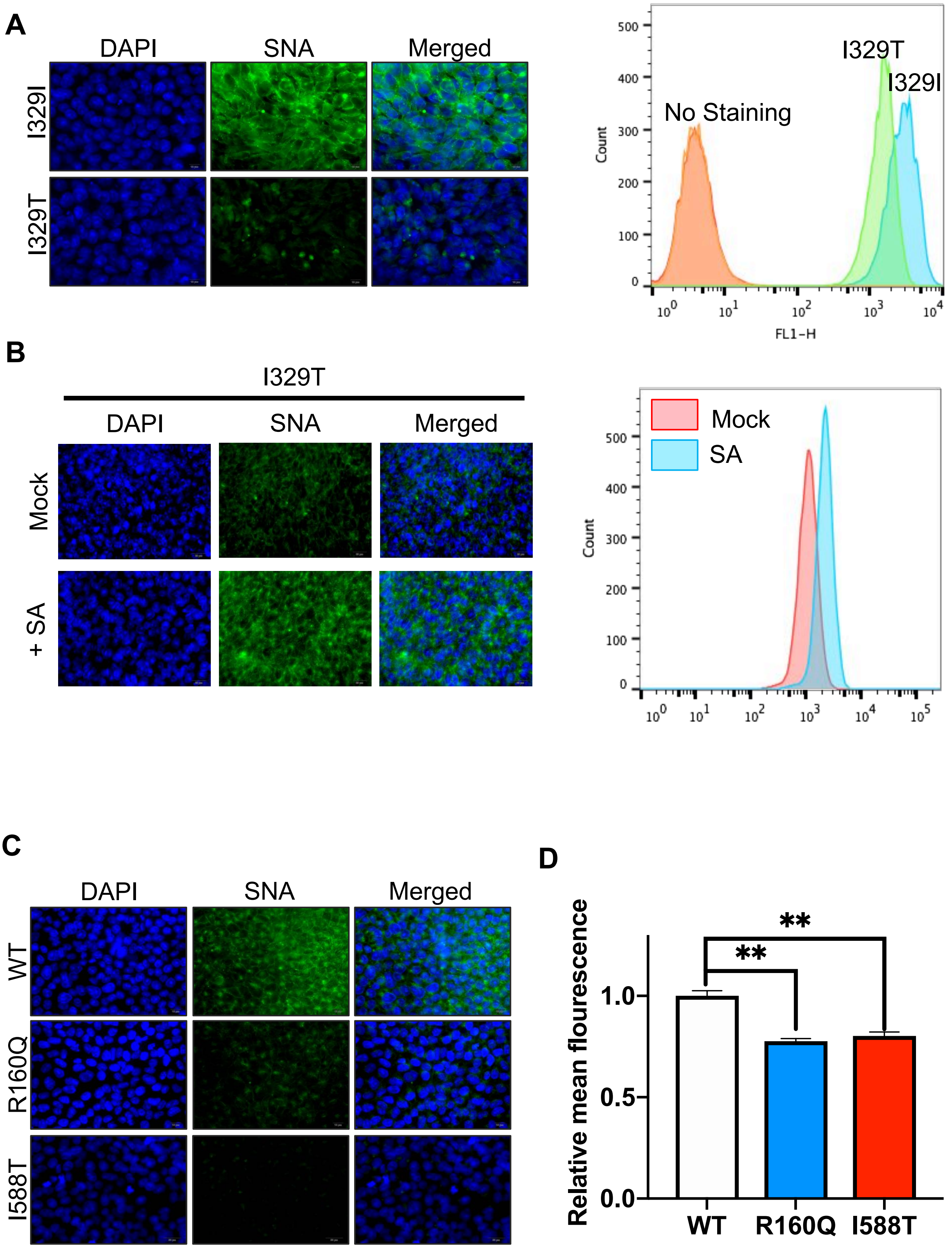
A
DAPI
SNA
Merged
I329I
I329T
No Staining
I329T
I329I
Count
FL1-H
B
I329T
DAPI
SNA
Merged
Mock
+ SA
Mock
SA
Count
C
DAPI
SNA
Merged
WT
R160Q
I588T
D
Relative mean flourescence
**
**
WT
R160Q
I588T

#### Figure. S5

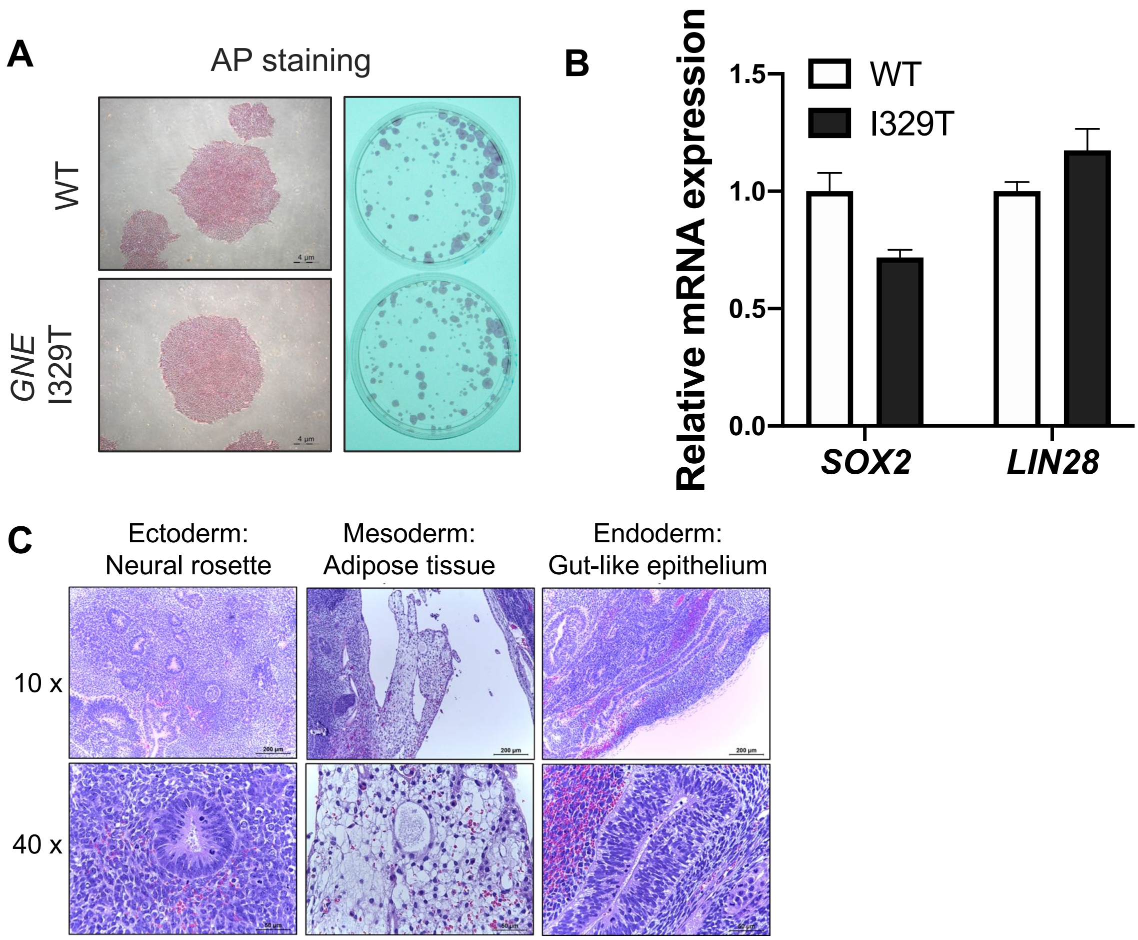

A
AP staining
WT
GNE I329T
4 µm
B
Relative mRNA expression
1.5
1.0
0.5
0.0
WT
I329T
SOX2
LIN28
C
Ectoderm: Neural rosette
Mesoderm: Adipose tissue
Endoderm: Gut-like epithelium
10 x
40 x
200 µm

### Figure. S6

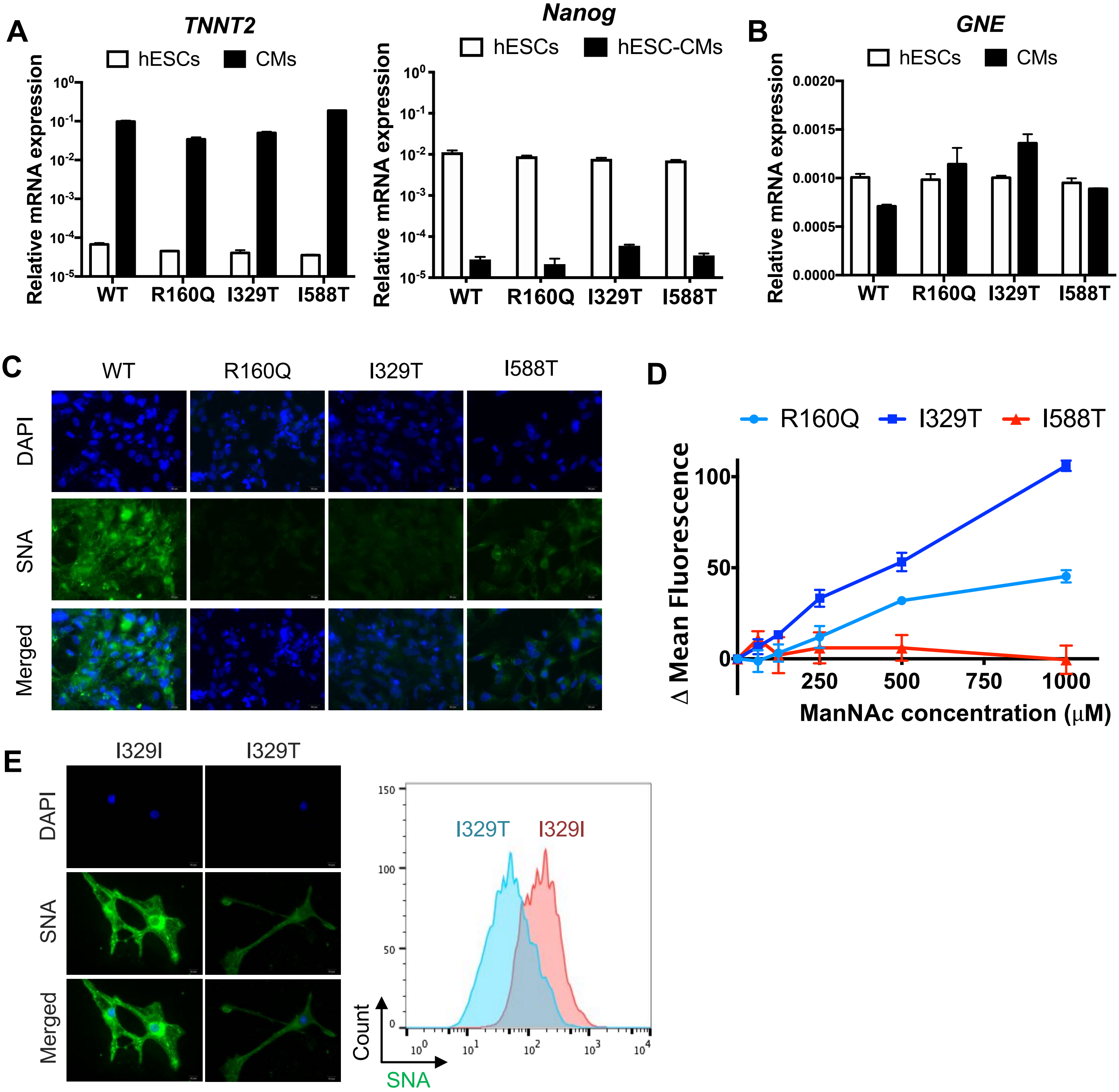

A
TNNT2
hESCs
CMs
Relative mRNA expression
WT
R160Q
I329T
I588T
Nanog
hESC-CMs
B
GNE
C
DAPI
SNA
Merged
D
Δ Mean Fluorescence
ManNAc concentration (μM)
E
I329I
Count

#### Table. S1

| Nucleotide change | Protein change NP_005467.1 (hGNE1) | Mutation type | Editing tool (Wild-type to mutant) | Editing tool (Disease to normal) | Clinical significance |
|---|---|---|---|---|---|
| c.893T>C | I298T | Transition | NGG-ABE | NGG-CBE | LP |
| c.1523T>C | L508S | Transition | NGG-ABE | NGG-CBE | LP |
| c.1556A>G | N519S | Transition | NGG-ABE | No PAM | LP |
| c.647T>C | V216A | Transition | NG-ABE | NG-CBE | P/LP |
| c.511A>G | M171V | Transition | NG-ABE | NG-CBE | P |
| c.1262T>C | V421A | Transition | NG-ABE | NG-CBE | LP |
| c.2135T>C | M712T | Transition | NG-ABE | No PAM | P |
| c.386G>A | R129Q | Transition | NGG-CBE | NGG-ABE | P/LP |
| c.1525C>T | H509Y | Transition | NGG-CBE | NGG-ABE | LP |
| c.1571C>T | A524V | Transition | NGG-CBE | NG-ABE | P/LP |
| c.1891G>A | A631T | Transition | NGG-CBE | NG-ABE | P |
| c.856C>T | Q286* | Transition | NGG-CBE | NG-ABE | LP |
| c.385C>T | R129* | Transition | NGG-CBE | NG-ABE | LP |
| c.529C>T | R177C | Transition | NGG-CBE | No PAM | LP |
| c.1892C>T | A631V | Transition | NG-CBE | NGG-ABE | P |
| c.1485G>A | W495* | Transition | NG-CBE | NGG-ABE | LP |
| c.-6C>T | Q30* | Transition | NG-CBE | NG-ABE | LP |
| c.484C>T | R162C | Transition | NG-CBE | NG-ABE | P/LP |
| c.829C>T | R277C | Transition | NG-CBE | NG-ABE | P/LP |
| c.175C>T | R59* | Transition | NG-CBE | NG-ABE | P |
| c.1258C>T | R420* | Transition | NG-CBE | NG-ABE | P |
| c.736C>T | R246W | Transition | NG-CBE | NG-ABE | P |
| c.22C>T | R8* | Transition | NG-CBE | No PAM | P |
| c.737G>A | R246Q | Transition | NG-CBE | No PAM | P/LP |
| c.655C>T | Q219* | Transition | NG-CBE | No PAM | LP |
| c.2122G>A | G708S | Transition | No PAM | NGG-ABE | LP |
| c.1727G>A | G576E | Transition | No PAM | NGG-ABE | P |
| c.1760T>C | I587T | Transition | No PAM | NGG-CBE | P/LP |
| c.916C>T | R306* | Transition | No PAM | NG-ABE | LP |
| c.2023T>C | Y675H | Transition | No PAM | NG-CBE | P |
| c.79C>T | P27S | Transition | No PAM | No PAM | LP |
| c.80C>T | P27L | Transition | No PAM | No PAM | LP |
| c.31C>T | R11W | Transition | No PAM | No PAM | LP |
| c.1306C>T | Q436* | Transition | No PAM | No PAM | LP |
| c.527A>T | D176V | Transversion | | | P/LP |
| c.4G>T | E2* | Transversion | | | P/LP |
| c.909T>A | C303* | Transversion | | | P |
| c.1844C>G | S615* | Transversion | | | P |
| c.38G>C | C13S | Transversion | | | P |
| c.1714G>C | V572L | Transversion | | | P |
| c.572C>G | S191* | Transversion | | | LP |
| c.445G>T | A149S | Transversion | | | LP |
| c.907_908delinsGT | C303V | Indel | | | P |
| c.1686del | C563fs | Deletion | | | P/LP |
| c.616+1del | | Deletion | | | P/LP |
| g.(?_36276884)_(36277052_?)del† | | Deletion | | | P |
| c.1546_1547del | | Deletion | | | P |
| c.640del | Y214fs | Deletion | | | LP |
| c.435_438del | I146fs | Deletion | | | LP |
| c.1376delG | | Deletion | | | LP |
| c.1740del | C581fs | Deletion | | | LP |
| c.1543_1544del | D515fs | Deletion | | | LP |
| c.1417del | S473fs | Deletion | | | LP |
| c.388del | I130fs | Deletion | | | LP |
| c.1411+1del | | Deletion | | | LP |
| c.1609_1616del | F537fs | Deletion | | | LP |
| c.1158del | K386fs | Deletion | | | LP |
| c.1781del | M594fs | Deletion | | | LP |
| g.(?_36227238)_(36227468_?)dup† | | Duplication | | | LP |
| c.636dup | | Duplication | | | LP |
| c.630dup | | Duplication | | | LP |
| c.680dup | | Duplication | | | LP |
| Single allele | | Copy number loss | | | P |

cDNA reference sequence: NM_005476.7
†Genomic DNA reference sequence: NC_000009.12
P, Pathogenic; LP, Likely pathogenic

**Table. S2**

| Cell | Mutation | Domain | Significance | Tool |
|---|---|---|---|---|
| H9 | R160Q | Epimerase | Pathogenic | NGG-CBE |
| | I329T | Epimerase | Likely pathogenic | NGG-ABE |
| | I588T | Kinase | Uncertain | NG-ABE |
| | V727M | Kinase | Conflicting | NGG-CBE |
| | V727M/I | Kinase | Unknown | NGG-CBE |
| | V727M, D726D/N | Kinase | Unknown | NGG-CBE |
| | V727M, D726N | Kinase | Unknown | NGG-CBE |
| | V727I, D726D/N | Kinase | Unknown | NG-CBE |
| | V727V/M | Kinase | Unknown | NG-CBE |
| | R160R/Q | Epimerase | Unknown | NG-CBE |
| CHA3 | I329T | Epimerase | Pathogenic | NG-ABE |
| | I329I | Epimerase | Silence mutation | NG-ABE |
| BJiPSC | R160Q | Epimerase | Pathogenic | NGG-CBE |
| | I588T | Kinase | Uncertain | NGG-ABE |
